# Exogenous antimicrobial straight-chain unsaturated fatty acids promote low-temperature growth of *Staphylococcus aureus*

**DOI:** 10.64898/2026.01.29.702679

**Authors:** Sharanya Paul, David Brewer, Matthew W. Frank, Arunachalam Muthaiyan, Vineet K. Singh, Antje Pokorny, Kelly M. Hines, Jan-Ulrik Dahl, Brian J. Wilkinson

## Abstract

The incorporation of oleic acid, a physiologically relevant and relatively non-toxic fatty acid, into membrane phospholipids and glycolipids has major impacts on the membrane biophysical properties of *Staphylococcus aureus*, including the promotion of low-temperature growth. Inclusion of oleic acid in growth media decreased the minimum growth temperature of *S. aureus.* It has not been studied yet whether other physiologically relevant, yet more antimicrobial straight-chain unsaturated fatty acids (SCUFAs), elicit similar responses. Here, we report that exogenous addition of these antimicrobial SCUFAs at 12°C led to their incorporation into the membrane lipids in only relatively small amounts, which, however, was sufficient to strongly promote growth at low temperatures. Intriguingly, these SCUFAs showed a preferential incorporation into diglucosyldiacylglyceride (DGDG) over phosphatidyl glycerol, and increased DGDG levels ∼ 4-fold in SCUFA-grown *S. aureus*. Two glycolipid-deficient mutants lost the growth-enhancing effects of SCUFAs at low temperatures and showed increased susceptibility to SCUFAs at 37°C, suggesting that glycolipids are important membrane components and are required for low-temperature adaptation. The presence of SCUFAs at low temperatures also enhanced the production of the carotenoid staphyloxanthin. Overall, our results suggest that multiple strategies are at play in the membrane lipid remodeling of *S. aureus* to adapt to the stress of low-temperature growth and likely to other environmental stresses. Incorporation of antimicrobial SCUFAs into membrane lipids may facilitate the insertion of free forms of these SCUFAs into the membrane and be a part of their mode of action.

## INTRODUCTION

*Staphylococcus aureus* is a major bacterial pathogen worldwide in both healthcare and community settings. The organism is versatile, adapting to multiple tissues and organs in which it may cause disease. Therapies for staphylococcal infections are challenging due to the emergence of methicillin-resistant *S. aureus*, which typically harbor resistant determinants to multiple antibiotics [1]. In addition, *S. aureus* is a leading cause of foodborne intoxications due to the proliferation of the organism in food and the release of enterotoxins [2].

Most, if not all, bacteria are able to incorporate exogenous fatty acids into membrane lipids to save carbon, energy, and reducing power over the *de novo* biosynthesis of endogenous fatty acids [3]. This can have profound effects on bacterial physiology, antimicrobial susceptibility, and may compromise susceptibility to bacterial type II fatty acid synthesis (FASII)-directed drugs by allowing bacteria to bypass the FASII pathway [4, 5]. Also, host fatty acids have been shown to exhibit major impacts on bacterial virulence [6]. When *S. aureus* is grown in conventional laboratory media, the biosynthesized membrane fatty acids incorporated into phospholipids and glycolipids are a mixture of branched-chain fatty acids and straight-chain fatty acids (BCFAs, SCFAs), respectively. Phosphatidylglycerol (PG) is the major *S. aureus* phospholipid species, and under these conditions, PG 32:0 with fatty acid *anteiso* C17:0 occupying the *sn-1* position, and *anteiso* C15:0 at the *sn-2* position is the major species [7]. *S. aureus* is believed to be unable to biosynthesize unsaturated fatty acids. The major straight-chain unsaturated fatty acid (SCUFA) in the animal body is oleic acid (C18:1Δ9), and PG 33:1, which contains C18:1 at the *sn-1* position and *anteiso* C15:0 at the *sn-2* position, is a major lipid species in cells grown *in vivo* in animal models [8]. It has been shown that *S. aureus* can incorporate substantial amounts of oleic acid from various sources, including the free fatty acid, glycerol trioleate, cholesteryl oleate, serum, and low-density lipoprotein particles via the fatty acid kinase and fatty acid binding protein FakAB system [9–13] (**FIG. 1**).

**FIG. 1:**
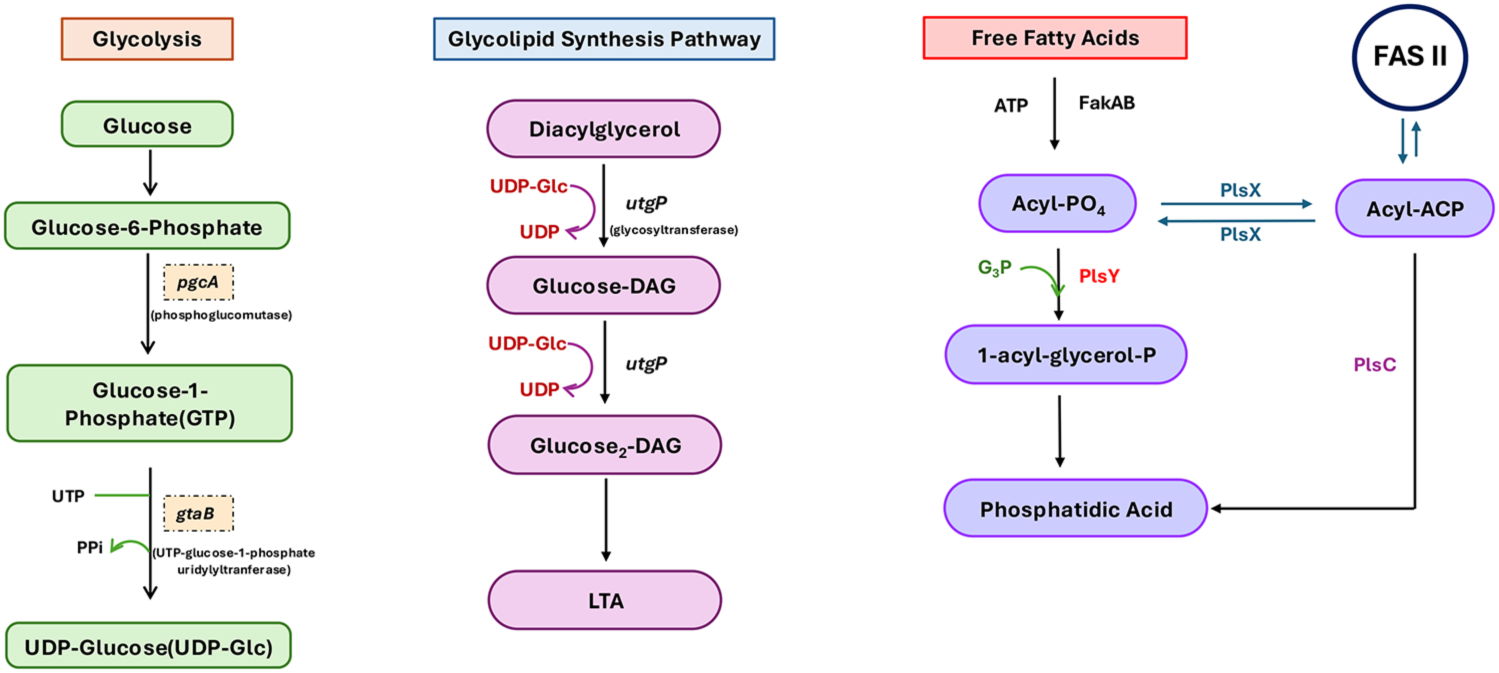
Biosynthesis of phospholipids, glycolipids, and LTA. Schematic representation shows the assimilation of free fatty acids via the FakAB system to form acyl-phosphate (Acyl-PO4), which is subsequently converted to acyl-ACP by PlsX for integration into the FASII system. Concurrently, the conversion of glucose-6-phosphate by the phosphoglucomutase *pgcA* drives glycolipid synthesis. Diacylglycerol (DG) and glucose-derived intermediates fuel LTA assembly, which is mediated by the glycosyltransferase *utgP*.

We have investigated the biochemistry, biophysics, and physiology of the incorporation of oleic acid into *S. aureus* membrane lipids. A detailed lipidomics investigation showed that oleic acid and its elongation product C20:1Δ11 were incorporated into both the *S. aureus* major lipids PG and diglucosyldiacylglyceride (DGDG) at the expense of PGs and DGDGs bearing endogenously biosynthesized BCFA and SCFA species [14]. Biophysical studies of lipid phase transitions in isolated membrane fragments, total lipid, phospholipid, and glycolipid fractions obtained from *S. aureus* grown in the presence and absence of oleic acid revealed that oleic acid incorporation into the cytoplasmic membrane led to increases in long-chain saturated and unsaturated glycolipid species that promote the formation of inverted lipid phases at higher temperatures [15]. Liposomes prepared from cells grown with oleic acid were less susceptible to permeabilization by the antimicrobial peptide Mastoparan X. We speculated that enrichment in unsaturated fatty acid species led to lipid packing defects and a more permeable cytoplasmic membrane that was counterbalanced by long-chain glycolipids. In physiological studies, we showed that the growth of *S. aureus* with 100 µM oleic acid resulted in SCUFAs becoming about 50% of the total fatty acids of the organism, that was accompanied by a marked stimulation of the rate of growth in cells subjected to the stress of low-temperature growth [14].

Given the significant biochemical, biophysical, and physiological consequences of incorporation of oleic acid by *S. aureus*, it is important to realize that there are other SCUFAs differing in chain length and degrees of saturation in the animal body. Several of these SCUFAs have much greater antimicrobial potency than oleic acid, and some are part of the innate immune system [16]. SCUFAs, to which *S. aureus* is likely to be exposed, are found in healthy human plasma in millimolar concentrations, including oleic acid (C18:1Δ9; 0.65-3.5 mM), sapienic acid (C16:1Δ6; 0.025-0.105 mM), palmitoleic acid (C16:1Δ9; 0.11-1.13 mM), linoleic acid (C18:2Δ9,12; 2.27-3.85 mM) and arachidonic acid (C20:4Δ5,8,11,14; 0.52-1.49 mM) [17]. Oleic acid and antimicrobial SCUFAs are also found esterified to glycerol and cholesterol in various tissues and fluids [10, 11].

Given the major impacts of the growth of *S. aureus* with oleic acid on the lipid composition of *S. aureus,* membrane biophysical properties, and adaptation to low temperatures, we felt that it was important to extend the studies to other SCUFAs. We hypothesized that these antimicrobial SCUFAs would also have major effects on membrane lipid biochemistry and on membrane functional properties, even though they are only tolerated at much lower concentrations than oleic acid. It is known that when SCUFAs are present at only low abundances in model liposomes they can have major impacts on membrane biophysical properties [18].

Although antimicrobial SCUFAs were only incorporated into low proportions of the total fatty acids when *S. aureus* was grown in sub-inhibitory concentrations of the fatty acids, they had a major impact on the lipidome and growth physiology of *S. aureus.* The SCUFAs increased DGDG levels and were preferentially incorporated into this molecule at 12°C. Studies of glycolipid-deficient mutants revealed that their growth at 12°C was non-responsive to SCUFAs, and that they were markedly more susceptible to the antimicrobial effects of SCUFAs at 37°C. This incorporation of SCUFAs and their elongation products into membrane phospholipids and glycolipids appeared to initiate a multicomponent lipid remodeling strategy to adapt to low temperatures involving changes in phospholipid and glycolipid species and enhanced production of staphyloxanthin. Incorporation of antimicrobial SCUFAs into membrane lipids may potentiate the antimicrobial activity of free forms of these SCUFAs. The results draw attention to the significant role of DGDG in the *S. aureus* membrane.

## RESULTS

### Sub-MIC concentrations of antimicrobial SCUFAs stimulate the growth of *S. aureus* at low temperatures

Barbarek et al. [14] showed that exogenous addition of 100 µM oleic acid in Tryptic Soy Broth (TSB) resulted in the incorporation of the SCUFA into phospho- and glyco-lipids, which sufficiently changed the *S. aureus* membrane composition and its physical properties to promote growth at 12°C. Since the minimum growth temperature for *S. aureus* is generally considered to be about 7°C in studies carried out using conventional bacteriological media [19], we were curious to know whether inclusion of 100 µM oleic acid into TSB would enhance the growth of the *S. aureus* strain JE2 on plates and in liquid cultures at temperatures lower than 12°C. After 30 days of growth at 10°C, 7°C and 4°C in TSB with or without oleic acid growth, on both plates and in liquid cultures was significantly enhanced in the presence of oleic acid **(SUPPLEMENTARY FIG. S1)**, indicating that SCUFAs may lower the minimum growth temperature of *S. aureus* and thus be significant to the food environment. Next, it was of interest to see whether other fatty acids could promote low-temperature growth and impact *S. aureus* membrane properties. The fatty acids initially studied were sapienic acid (SA, C16:1Δ6), palmitoleic acid (PA, C16:1Δ9), linoleic acid (LA, C18:2Δ9,12), and arachidonic acid (AA, C20:4Δ5,8,11,14). These SCUFAs are antimicrobial fatty acids with minimum inhibitory concentrations (MICs) of about 31 µM (*i.e.* SA, PA, and AA) and 250 µM (*i.e.* LA), respectively (**SUPPLEMENTARY Table S1**) [20]. In contrast, oleic acid has an MIC of 500 µM or more [20]. Accordingly, TSB supplemented with a range of SCUFAs in concentrations from 0.1-100 µM in 96-well plates was inoculated with *S. aureus* strain JE2 and cultivated for 5 days at 12°C. Twelve°C was chosen as the experimental temperature for practical reasons because the growth rate was slowed significantly yet experiments could be completed in a reasonable time frame as compared to 7°C (days vs weeks, see **SUPPLEMENTARY FIG. S1**). **FIG. 2** illustrates differences in the growth of JE2 in TSB supplemented with various concentrations of the indicated fatty acids at 12°C compared to the ethanol-treated controls. Growth was promoted by all concentrations of oleic acid tested up to 100 µM (**FIG. 2A**). In contrast, while all four antimicrobial SCUFAs tested also stimulated growth over the ethanol control at the lower concentrations tested, their growth stimulatory effect began to diminish after 30 µM, and higher concentrations led to growth inhibition, which is in stark contrast to oleic acid. This is in line with the MICs of these SCUFAs (about 30 µM), compared to oleic acid (> 500 µM) (**SUPPLEMENTARY Table S1**) [20].

**FIG. 2:**
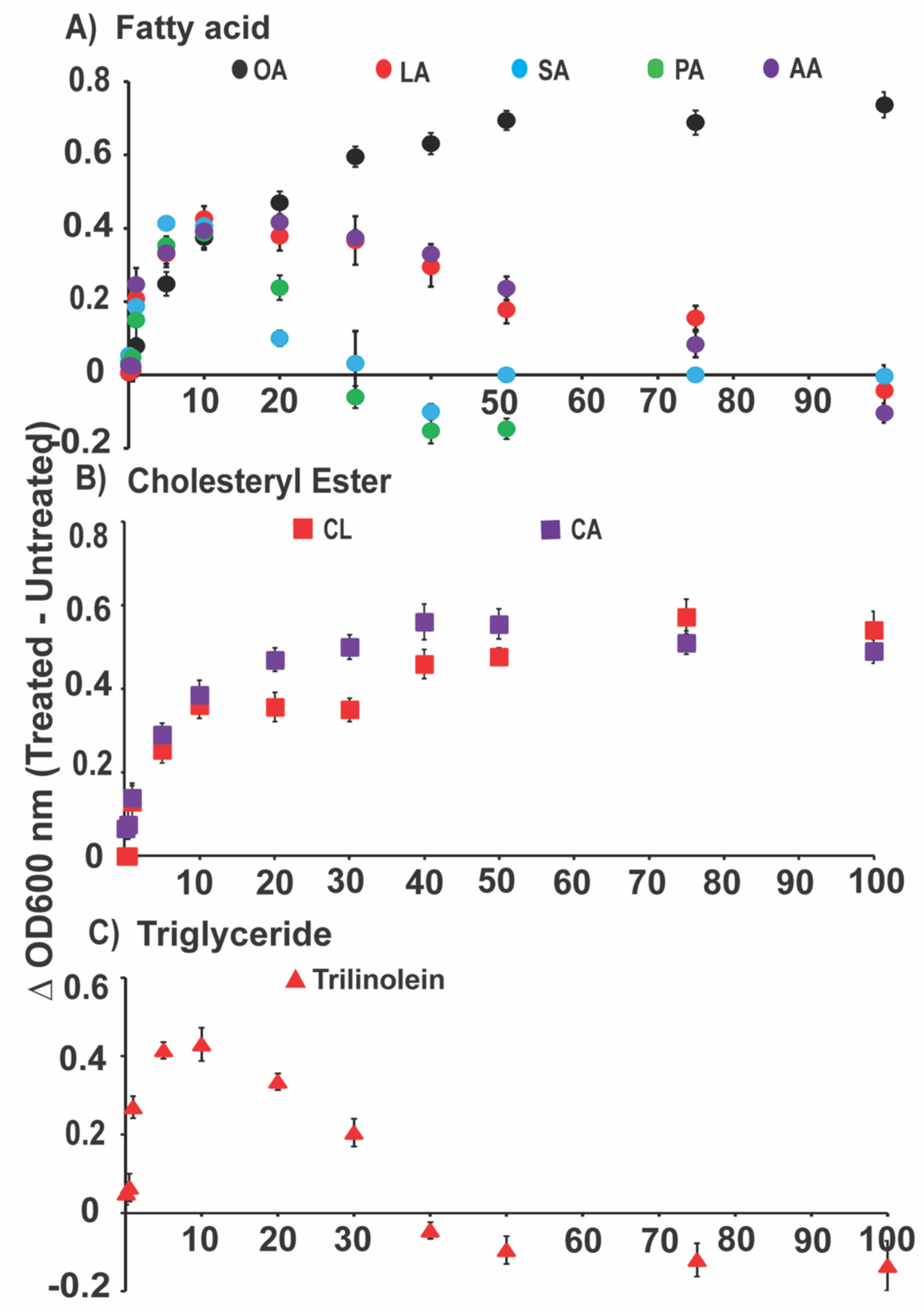
Free and esterified SCUFAs promote the growth of *S. aureus* strain JE2 at 12°C. Growth was assessed in 96-well plates by measuring OD_600_ after 5 days at 12°C. TSB was supplemented with varying concentrations (0.1–100 µM) of the indicated **(A)** unsaturated fatty acids (circles), **(B)** cholesteryl esters (squares), and **(C)** the triglyceride of linoleic acid, trilinolein (triangles). Growth was normalized by subtracting the OD_600_ of the ethanol-only control. **(A) Oleic acid (OA)** promoted growth at all tested concentrations up to 100 µM. Low concentrations (≤30 µM) of **sapienic (SA)**, **linoleic (LA)**, **palmitoleic (PA)** and **arachidonic acids (AA)** stimulated growth, but higher concentrations led to inhibition. **(B)** The cholesteryl esters **cholesteryl linoleate (CL)** and **cholesteryl arachidonate (CA)** exhibited a concentration-dependent effect on growth, with higher concentrations resulting in greater stimulation. **(C) Trilinolein** showed similar trends to free fatty acids: low concentrations stimulated growth, whereas higher concentrations were toxic, (n=3, ±S.D.).

### Stimulation of low temperature growth by glycerol and cholesterol esters of SCUFAs and serum

Hines et al. [10] have shown that triglycerides and cholesteryl esters of oleic acid and linoleic acid can act as donors for SCUFA incorporation into phospholipids and glycolipids following hydrolysis of the esters by the lipase Geh. An interesting observation was made when the low-temperature growth-promoting activity of cholesteryl esters was tested. The cholesteryl esters promoted low-temperature growth but were tolerated to significantly higher concentrations (**FIG. 2B**) than were the free fatty acids or their triglycerides (**FIG. 2A&C)**. The tolerance of cholesteryl esters to higher concentrations by *S. aureus* suggests that cholesterol may have a protective effect on the membrane against these antimicrobial SCUFAs. To assess whether cholesterol protects *S. aureus* from SCUFA-mediated toxicity at 12°C, we exposed JE2 to free linoleic acid, its cholesteryl ester cholesteryl linoleate, and a combination of linoleic acid and exogenous cholesterol at 300 μM and monitored growth for 5 days. Addition of linoleic acid significantly diminished the growth stimulatory effects when supplemented at concentrations higher than 20 μM, while addition of the cholesteryl ester counterpart promoted growth at all concentrations tested up to 100 µM (**Fig. 3A**). Strikingly, the addition of exogenous cholesterol along with linoleic acid resulted in full growth recovery, mimicking the effect of cholesteryl linoleate and suggesting a protective, possibly membrane-stabilizing role for cholesterol. We observed a very similar outcome with free arachidonic acid, its cholesteryl ester cholesteryl arachidonate, and a combination of arachidonic acid and exogenous cholesterol (**SUPPLEMENTARY FIG. S2A**). The addition of cholesterol alone had no effect on *S. aureus* growth. Triglycerides of various SCUFAs were shown to enhance growth at a slightly lower range of concentrations than the free fatty acids (**FIG. 2C**), presumably due to the hydrolysis of triglyceride by the lipase Geh, yielding free SCUFAs that are incorporated into lipids and promote growth [21]. In support of this, there was no promotion of growth at 12°C of a *geh* mutant by cholesteryl linoleate (**FIG. 3B**). Heat-treated human serum at a concentration of 20%, known for its ability to supply SCUFAs and cholesterol to *S. aureus* [10], also stimulated growth at 12°C (**SUPPLEMENTARY FIG. S2B**).

**FIG. 3:**
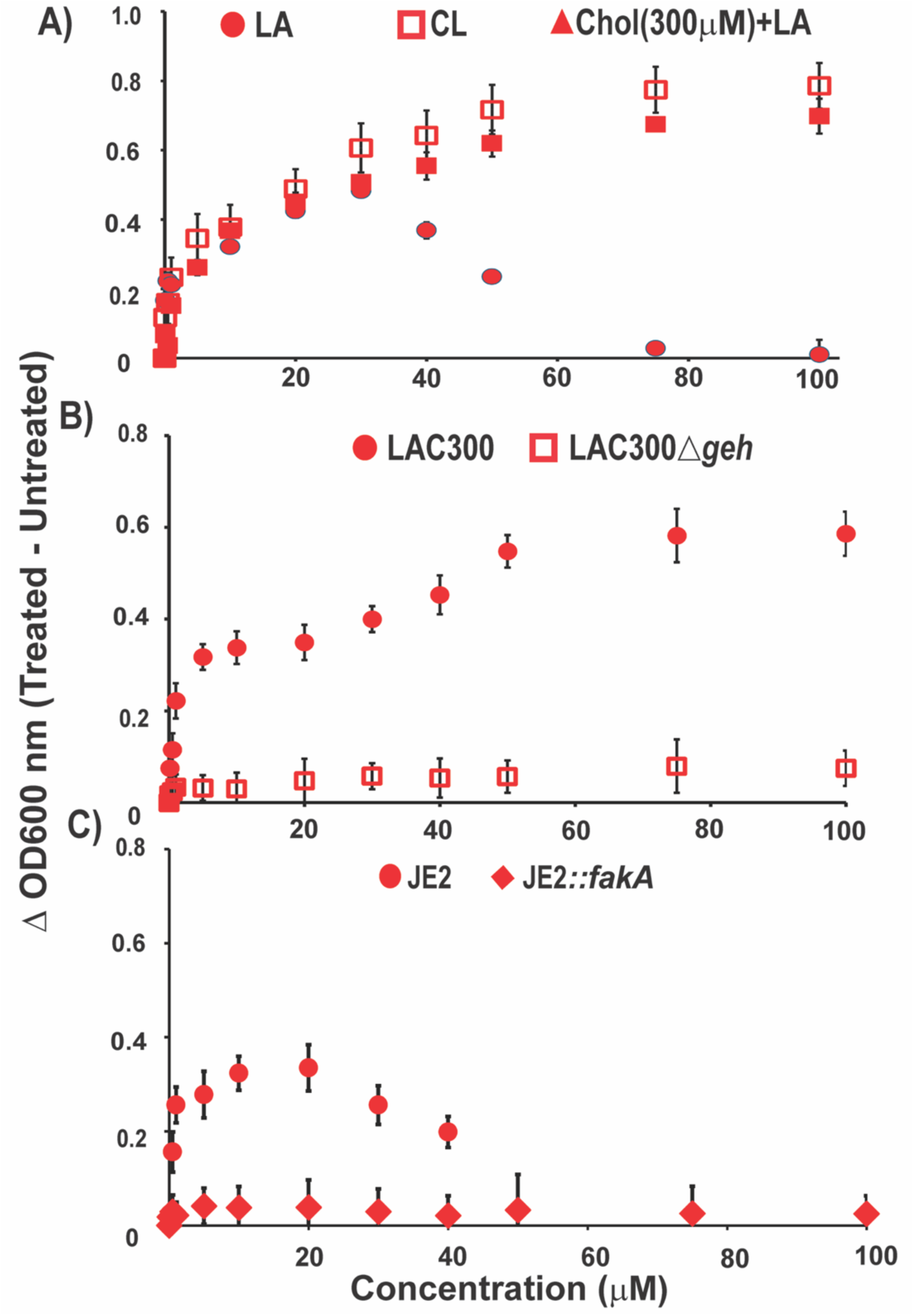
Growth stimulation of *S. aureus* at 12°C is dependent on FakA and maintained at higher concentrations when fatty acids are esterified with cholesterol. **(A)** Overnight cultures of JE2 were diluted into fresh TSB to an OD_600_ ∼0.05 and cultivated in 96-well plates at 12°C in the presence and absence of the indicated concentrations of linoleic acid (LA, filled circle), cholesteryl linoleate (CL, unfilled squares), and co-incubation of linoleic acid with 300 µM cholesterol (LA+ Chol, filled triangles), respectively. OD_600_ was determined after 5 days and normalized by subtracting the OD_600_ of the ethanol-only control in case of LA and CL. And for the co-incubation (Chol+LA) OD_600_ was determined after 5 days and normalized by subtracting the OD_600_ of the cholesterol-only control. Free LA promotes growth, peaking around 20 to 30 µM, but is inhibitory at higher concentrations. In contrast, growth stimulation by CL or the exogenous addition of 300 µM cholesterol to LA is sustained at all concentrations tested, (n=3, ± SD). **(B)** LAC300 (circles) and LAC300*Δgeh* (squares) were grown in TSB media in the presence and absence of the indicated concentrations of CL in 96-well plates at 12°C. OD_600_ was determined after 5 days and normalized by subtracting the OD_600_ of the ethanol-only control. CL enhances growth at 12°C in a *geh*-dependent manner, (n=3, ± SD). **(C)** JE2 (circles) and JE2::*fakA* (diamonds) were grown in TSB media in the presence and absence of the indicated concentrations of LA for 5 days at 12 °C. OD_600_ was determined after 5 days and normalized by subtracting the OD_600_ of the ethanol-only control, (n=3, ±S.D.).

### The stimulatory effect of SCUFAs on low temperature growth is dependent on incorporation into lipids via the FakAB system

The effects of linoleic acid and arachidonic acid on the growth of the FakA-defective mutant at 12°C were assessed. Compared to the parent strain, little growth occurred in the JE2 *fakA::Tn* mutant in the presence of linoleic acid (**FIG. 3C**) or arachidonic acid **(SUPPLEMENTARY FIG. S2C)** and growth resembled that of the ethanol control culture. These findings indicate that growth stimulation at low temperatures is dependent upon covalent incorporation of SCUFAs into phospho- or glyco-lipid structures (**FIG. 1**), and simple insertion of SCUFA into the membrane as a free fatty acid is ineffective in promoting low-temperature growth.

### Carotenoid production accompanies the growth stimulatory effects of SCUFAs at 12°C

Barbarek et al. [14] noted increased staphyloxanthin production in cells grown at 12°C in TSB supplemented with oleic acid. It was of interest to see whether this extended to other SCUFAs. Fold-changes in carotenoid production at 12°C over 37°C were determined for *S. aureus* cultivated in the presence and absence of 10 μM oleic acid, sapienic acid, linoleic acid, arachidonic acid, and palmitoleic acid, respectively. Ten µM concentrations of these SCUFAs caused marked 4-to 7-fold stimulation of staphyloxanthin production in the wildtype (WT) strain JE2 at 12 °C compared to 37 °C (**FIG. 4A**). As expected, no such stimulatory effect was observed in the ethanol controls or in the JE2 (*crtM::Cm^R^)* strain, which is deficient in staphyloxanthin production [22]. Staphyloxanthin production was restored in the carotenoid-deficient mutant complemented with the pCU*crtOPQMN* plasmid expressing the carotenoid biosynthesis operon (**FIG. 4A**). In the presence of oleic acid and arachidonic acid, the complemented strain produced significantly more carotenoid than the parent strain. Next, the impact of staphyloxanthin induction by these various fatty acids on the growth of parent strain JE2, the carotenoid-deficient mutant (*crtM::Cm^R^)* and the complemented strain (*crtM::Cm^R^*+pCU*crtOPQMN*) at 12°C was assessed (**FIG. 4B-E**). In all cases, the carotenoid-deficient mutant grew less well than the carotenoid-producing parent strain but significantly better than the ethanol control. Interestingly, complementation with the pCU*crtOPQMN* plasmid was only partially successful. In light of a similar growth response of the complemented strain compared to the carotenoid-deficient strain, it could be that enhanced production of staphyloxanthin in the complemented strain occurs in excess of what is required and can be tolerated for optimal cold adaptation. However, our results indicate that staphyloxanthin production is significantly enhanced by the combination of low temperature and SCUFAs at low temperatures, and may play a significant role in cold adaptation, but is not absolutely required.

**FIG. 4:**
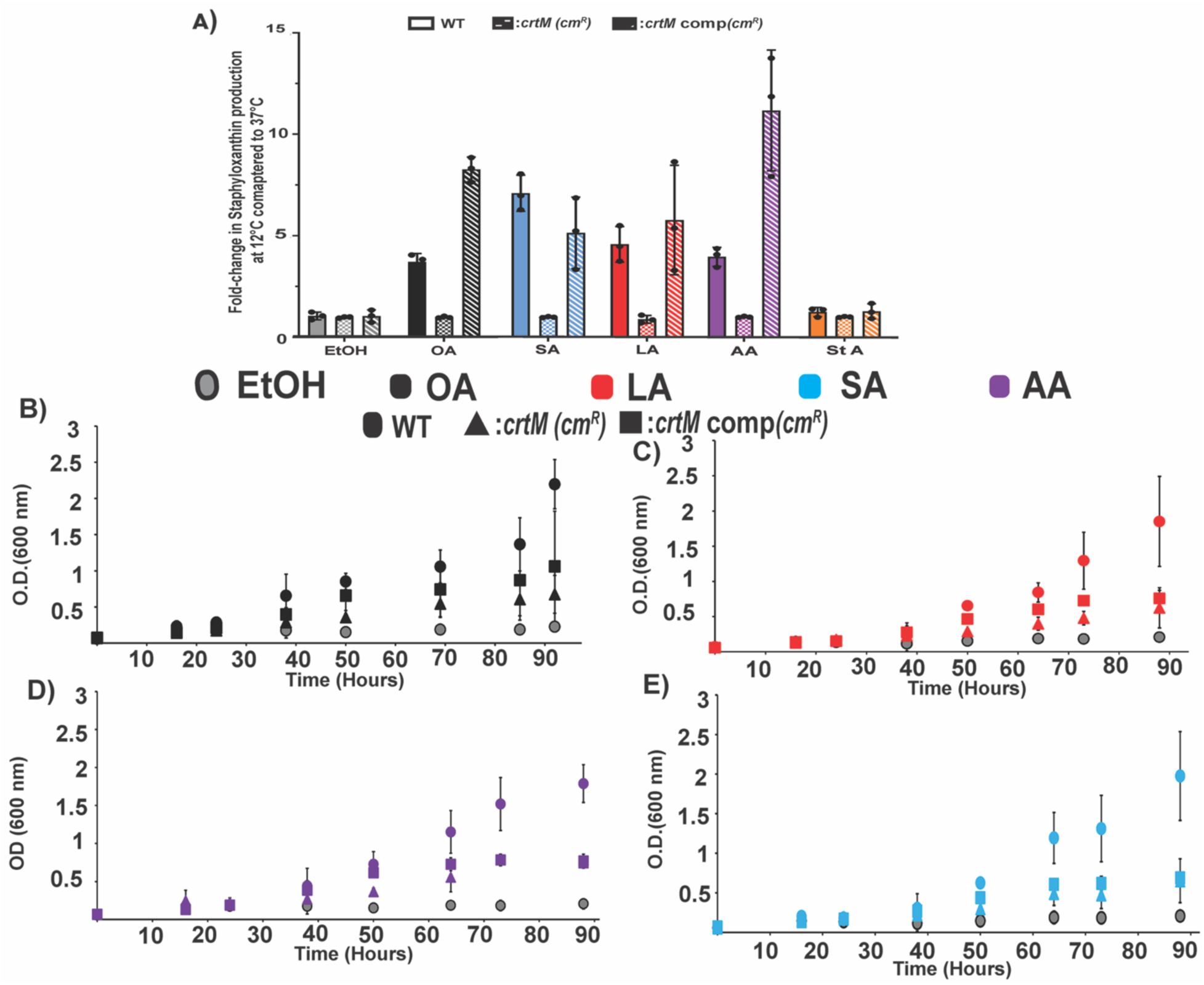
Staphyloxanthin production, enhanced by SCUFAs during growth at 12 °C, contributes to the cold adaptation of *S. aureus*. **(A)** *S. aureus* JE2 was grown for 5 days at 12°C in the presence and absence of 10 µM of **oleic acid (OA)**, **linoleic acid (LA)**, **arachidonic acid (AA)**, **sapienic acid (SA)** and **steric acid (StA).** Cells were washed twice in phosphate-buffered saline (PBS) and normalized to 5 mL of an O.D._600 nm_ = 2. Staphyloxanthin was extracted with warm methanol (55°C) for 5 min, and the O.D._465 nm_ of the supernatant was determined, (n=3, ±S.D.).**(B-E)** Growth of the *S. aureus* wild-type strain JE2 (circles), the carotenoid-deficient mutant (*crtM::Cm^R^*) (triangles), and the complemented strain ( *crtM::Cm^R^*+pCU*crtOPQMN*) (squares) was recorded at 12 °C in the presence of 10 μM (**B**) **(OA)**, (**C**) **(LA)**, (**D**) **(AA)**, and (**E**) **sapienic acid (SA)** at the indicated time points. Ethanol-treated cultures served as controls. JE2 showed enhanced growth with all fatty acids, whereas the *crtM::Cm^R^* strain showed reduced growth but still outperformed the ethanol-only control. The complemented strain (*crtM::Cm^R^*+pCU*crtOPQMN*) did not fully restore the wild-type phenotype, (n=3, ±S.D.).

### Survey of the effects of a range of SCUFAs on the promotion of growth at low temperatures

In order to evaluate the impact of carbon chain length and number of double bonds on the stimulation of low-temperature growth to a wider range of SCUFAs in a high throughput manner, the growth of JE2 cultures supplemented with 10 and 100 μM concentrations of various SCUFAs was monitored. All the SCUFAs tested stimulated the growth of the strain at 12°C at low temperature at concentrations of 10 µM, with the exception of *trans*-vaccenic acid (C18:1Δ11 *trans*) (**Supplementary Table S1**). However, at 100 µM concentrations, the most growth stimulatory SCUFAs were various forms of C18:1, *i.e.* oleic acid (C18:1Δ9 *cis*), elaidic acid (C18:1Δ9*trans*), *cis*-vaccenic acid (C18:1Δ11*cis*) and *trans*-vaccenic acid (C18:1Δ11 *trans*) (**Supplementary Table S1**). All the polyunsaturated fatty acids tested were inactive in growth promotion at 12°C at concentrations of 100 µM, likely due to toxicity. Initially, it was surprising that the two *trans* isomers promoted low-temperature growth given the slight kink only in the molecule imparted by the *trans* double bond. Accordingly, the impact of 100 μM concentrations of elaidic acid with the double bond in the *trans* configuration (*i.e.,* EA), *cis*-vaccenic acid (*i.e., cis*-VA), and the *trans*-form of vaccenic acid (*i.e., trans-VA*) on growth at 12°C in shaking flasks was determined. We confirmed that all three stimulated cold temperature growth, although *trans*-VA was somewhat less effective than the other fatty acids (**FIG. 5**). Inclusion of any of these fatty acids in the medium also enhanced carotenoid production. In contrast, the fully saturated C18:0 stearic acid (*i.e.,* StA) neither showed growth promoting activity nor stimulated staphyloxanthin production (**FIG. 4A**, **FIG. 5**).

**FIG. 5:**
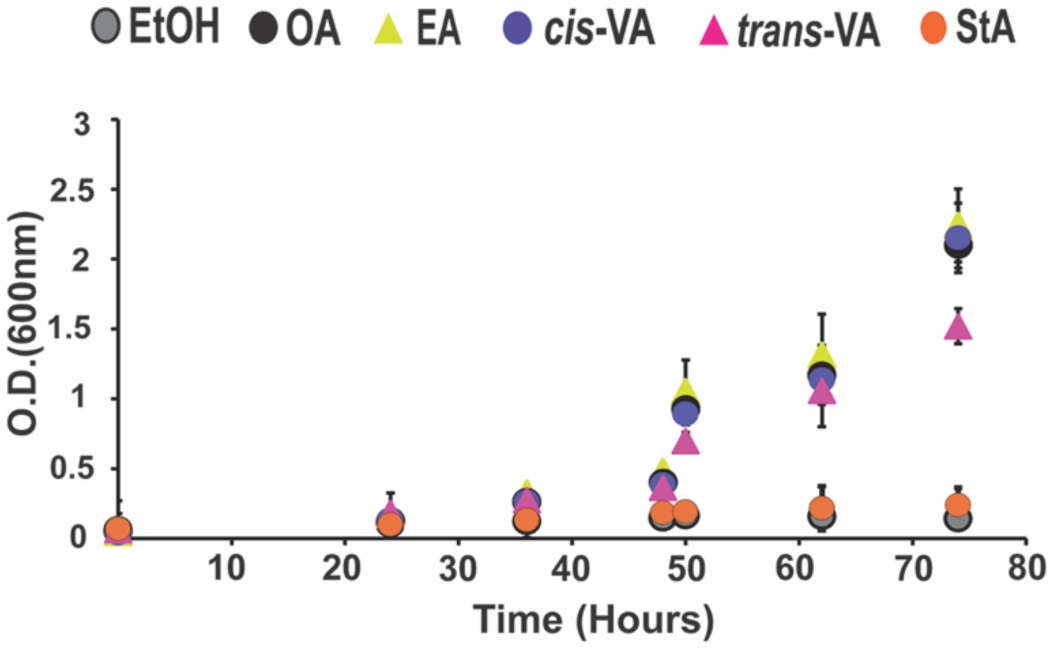
*cis* and *trans* isomers of C18:1Δ9 and C18:1Δ11 promote growth at 12°C. Cultures supplemented with **elaidic acid (EA, trans-18:1Δ9)**, **cis-vaccenic acid (cis-VA, cis-18:1Δ11)**, or **oleic acid (OA, cis-18:1Δ9)** exhibited enhanced growth comparable to each other, while **trans-vaccenic acid (trans-VA, trans-18:1Δ11)** provided a moderate growth benefit. No growth stimulation was observed in the presence of **stearic acid (StA)**, (n=4, ±S.D.).

### Lipidomics characterization of the growth adaptation with subinhibitory antimicrobial SCUFAs at 12°C

Of particular interest for us was what was the impact of those antimicrobial SCUFAs on the lipidome of *S. aureus*, as this may reveal important information to explain the observed growth stimulatory effects. When the SCUFAs were present in cultures at 37°C and 12°C, they were incorporated into both *S. aureus* major lipid species PG and DGDG, yielding modified unique PG and DGDG species that set them apart from cells grown in unsupplemented TSB medium (**FIG. 6-7; Supplementary FIG. S3-4**). While monounsaturated SCUFAs and linoleic acid did not impact saturated PG levels, arachidonic acid explicitly suppressed shorter saturated chains like PG 30:0 and PG 31:0 at 37°C (**FIG. 6**). The incorporation of these exogenous fatty acids shifted the unsaturated PG profile significantly, showing a strong preference for adding FA 18:1 over FA 16:1 into PG acyl tails (**FIG. 6**). Linoleic acid produced its own major unsaturated PG variants, while arachidonic acid predominantly yielded PG 35:4 at 37°C. Supplementation with exogenous SCUFAs alters the DGDG lipid population at 37°C by reducing certain saturated species and incorporating various fatty acids into new DGDG species, mirroring largely those observed in unsaturated PG populations. A more detailed description of how supplementation with antimicrobial SCUFAs affects lipid profiles at 37°C is provided in the *Supplementary Material*.

**FIG. 6:**
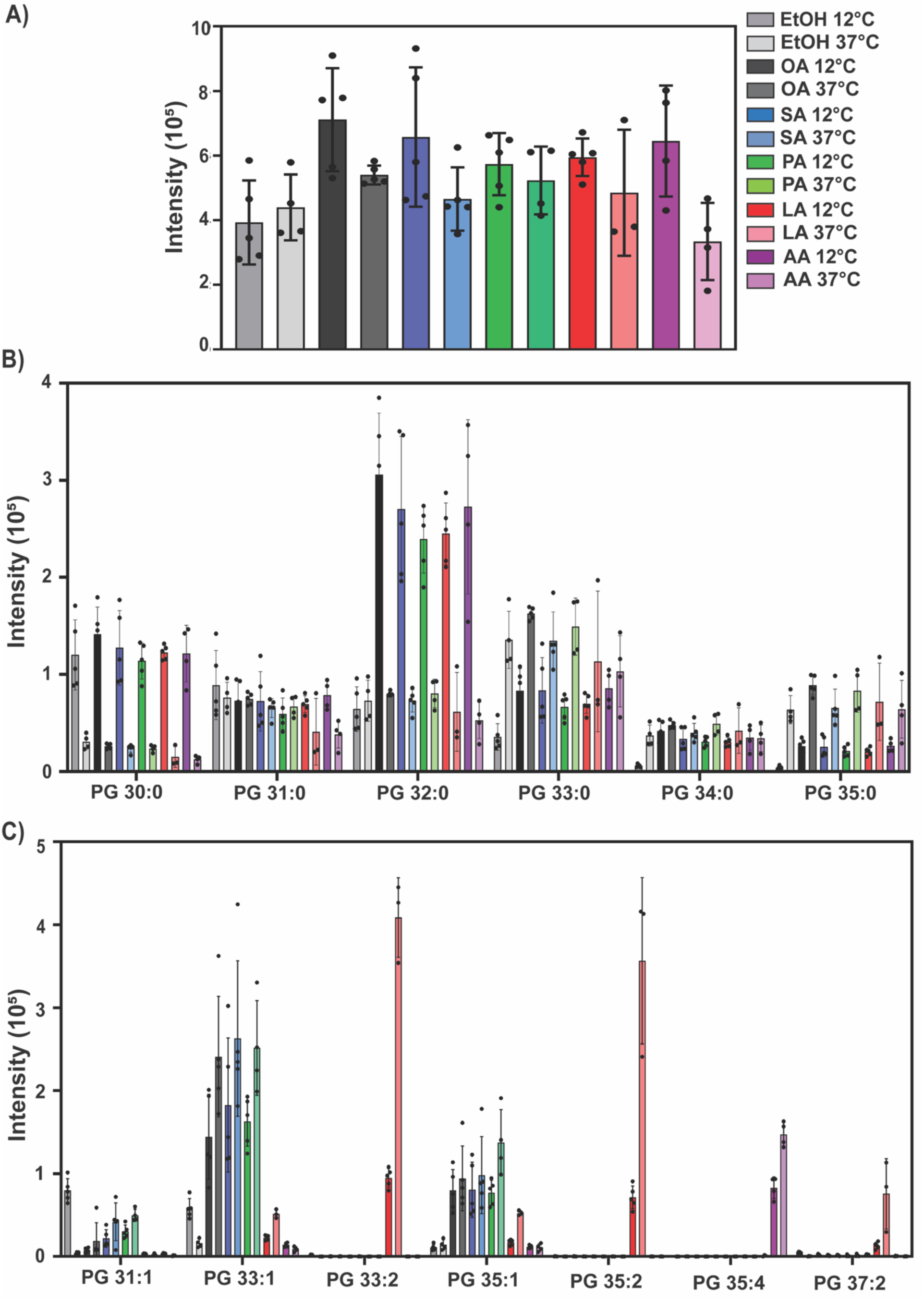
Lipidomics analysis of PG lipids in JE2 grown at 12°C and 37°C grown with varying exogenous free fatty acids at 10 μM. **(A)** Total PG abundance across all species. **(B)** Impact on native saturated PG species. **(C)** High-abundance unsaturated PG species, (n=5, ±S.D.).

Low-temperature growth at 12°C promotes a shift toward PG species with shorter acyl tails and a decrease in long-chain acyl tails, a trend largely independent of SCUFA presence (**FIG. 6**). While the incorporation of exogenous SCUFAs into staphylococcal PGs is significantly greater at 37°C than at 12°C, the overall amount of these unsaturated PGs remains nearly 100-fold lower than endogenous saturated species. In the absence of SCUFAs, low temperature growth had no impact on total PG levels for the EtOH control (**FIG. 6A**). When SCUFAs were provided, a slight but statistically insignificant increase in total PGs was observed in bacteria grown at 12°C. However, the analysis of PGs at the level of individual lipid species revealed greater variation in the abundance and structure of these lipids under the conditions of low temperature and SCUFA supplementation (**FIG. 6**; *see Supplementary Material*). The most dramatic differences between the membrane lipids of *S. aureus* grown at 12°C versus 37°C was observed in the DGDG lipid class. The total abundance of DGDGs was nearly 400% higher in bacteria grown at 12°C than at 37°C when SCUFAs were provided in the growth media (**FIG. 7**). Underlying this global impact was an increase in endogenous DGDGs with short acyl tails in nearly all bacteria grown at 12°C, which mirrored a similar trend observed for PGs. However, the increase in short-tailed DGDGs did not occur with a concomitant decrease in long-tail species like DGDG 35:0 (**FIG. 7B**). In contrast to PGs, the incorporation of SCUFAs into staphylococcal DGDGs occurred to higher levels and unsaturated DGDG species were 1/10^th^ the abundance of saturated species (versus 1/100^th^ for PGs). The preferences for incorporation of the various SCUFAs was more pronounced within the DGDG species, as well. The short-tailed SCUFAs, derived from sapienic acid and palmitoleic acid, were esterified into DGDG 33:1 at higher amounts than the other monounsaturated fatty acid, oleic acid, but there was no difference in abundance as a function of growth temperature for any of the three monounsaturated fatty acids. DGDG 35:1, on the other hand, was higher in abundance at 12°C for all three monounsaturated fatty acids, with the largest fold-change increase from the incorporation of oleate at 12°C (**FIG. 7**). As observed for PGs, the use of linoleate to biosynthesize poly-unsaturated DGDGs was greater at 37°C than 12°C. However, the fold-change difference between the two growth temperatures was smaller for linoleate-containing DGDGs than linoleate-containing PGs. Although arachidonate was minimally incorporated into PGs, DGDGs containing arachidonate-derived acyl tails were present at high abundance at both growth temperatures, with no significant difference, when arachidonic acid was provided in the medium. Low levels of DGDGs containing two exogenously derived SCUFA acyl tails (e.g., DGDG 34:2 and 36:2) were also detected in the bacteria that were provided with monounsaturated fatty acids. Trace amounts of double linoleate incorporation (e.g., DGDG 38:4) was detected, as well **(Supplementary FIG. S3)**. In contrast to PGs and DGDGs, where total amounts of these lipids were elevated in the 12°C growth conditions only in the presence of SCUFAs, we detected an increase in the amount of LysylPGs (**SUPPLEMENTARY FIG. S3-4**) in bacteria grown at 12°C independent of SCUFA supplementation. The magnitude of the increase did vary between the broth supplementation conditions, though. A 2-fold rise in LysylPG levels was observed between the 12°C and 37°C ethanol control conditions, whereas the difference between LysylPG level at 12°C and 37°C was greater than 4-fold when arachidonic acid was provided during the growth. At the level of individual species, LysylPGs followed similar trends to their biosynthetic precursor, PGs. Compared to previous studies in cells grown with higher concentrations of oleic acid (*i.e.,* 100 µM), where the percentage of unsaturated species were about 50% [7, 14, 15], sapienic acid, palmitoleic acid, oleic acid, and arachidonic acid at 10µM were only incorporated to between 6 to 11%, yet this was sufficient to elicit a comparable growth stimulation (**FIG. 2**).

**FIG. 7:**
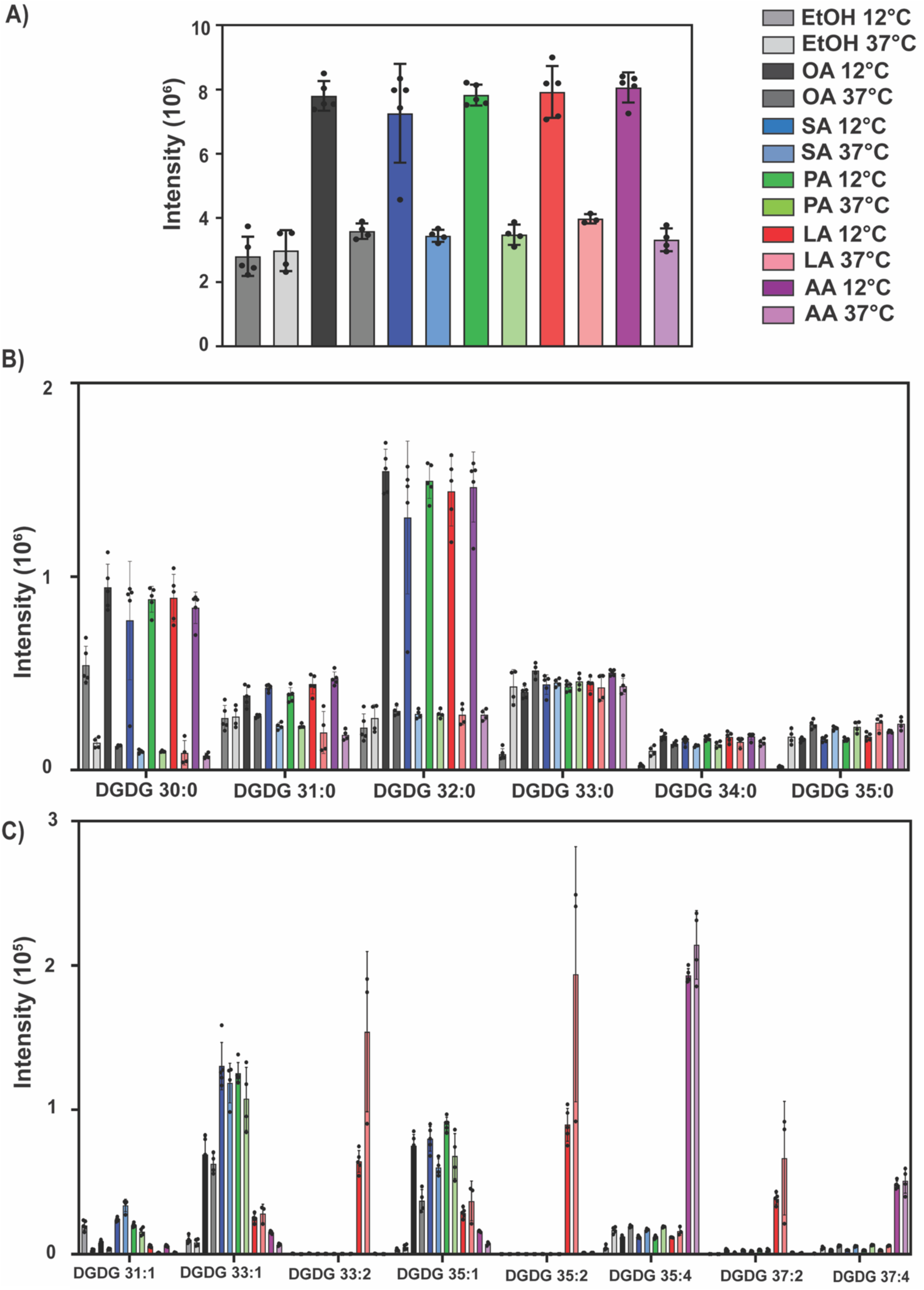
Lipidomics analysis of DGDG lipids in JE2 grown at 12°C and 37°C. grown with varying exogenous free fatty acids at 10 μM. **(A)** Total DGDG abundance across all species. **(B)** Impact on Native saturated DGDG species. **(C)** High-abundance unsaturated DGDG species, (n=5, ±S.D.).

### Lipidomic impact of C18:1 Fatty Acids

Amongst the DGDGs, low-temperature growth with ethanol as the control led to a greater increase in endogenous short-chained species **(SUPPLEMENTARY FIG. S5)** than was observed for PGs from the same bacteria or in the previous experiment (**FIG. 7**). As above, the incorporation of SCUFAs was far more abundant in DGDGs than PGs. SCUFAs with *trans*-geometry double bonds were incorporated into DGDGs to a greater extent at 12°C than at 37°C, whereas the *cis*-geometry SCUFAs were incorporated into DGDGs to similar degrees at both growth temperatures **(SUPPLEMENTARY FIG. S5)**. Again, the incorporation of exogenous 18-carbon SCUFAs onto both glycerol backbone positions was more prevalent at the low growth temperature with the highest incorporation for *cis*-vaccenic acid **(SUPPLEMENTARY FIG. S5)**. DGDGs containing the products for FASII elongation of the 18-carbon SCUFAs, DGDG 38:2 and 40:2, were significantly higher in abundance for the *cis*-geometry isomers oleic acid and *cis*-vaccenic acid at 12°C relative to 37°C and the *trans*-isomers. The various C18:1 fatty acids showed similar impacts on the LysylPG species to their impact on the PG species **(SUPPLEMENTARY FIG. S6)**.

### Glycolipid-deficient mutants did not show stimulation of growth at 12°C by SCUFAs and were more susceptible to SCUFAs at 37°C

Due to the substantially SCUFA-induced increase in DGDG levels observed in the lipidome of *S. aureus* grown at 12°C, we monitored the responses of two glycolipid-deficient mutants with transposon insertions in *gtaB* (UTP-glucose-1-phosphate uridylyltranferase) and *ugtP* (glycosyltransferase) (**FIG. 1**) to 10 and 100 μM oleic acid and 10 μM of sapienic acid at 12°C in flask cultures (**FIG. 8**). The results show a much-diminished growth stimulation response to SCUFAs at 12°C in the *gtaB* and *ugtP* mutants compared to the parent strain indicating that glycolipids are critically required for adaptation to low temperature. At 37°C the two glycolipid-deficient mutants showed a markedly different response to the parent strain JE2 to SCUFAs. Strain JE2 did not show inhibition by 10 µM sapienic acid or 10 and 100 µM oleic acid, although 100 µM oleic acid caused some initial growth delay (**FIG. 9A**). In contrast, both glycolipid-deficient mutants were completely susceptible to 100 µM oleic acid, with growth being entirely inhibited under these conditions (**FIG. 9B and C**). Furthermore, 10 µM sapienic acid was considerably more inhibitory to the mutant strains than to JE2, indicating that the absence of glycolipids increases susceptibility to antimicrobial fatty acids. Lipidomic analyses of the two mutants confirmed the absence of DGDGs in these strains.

**FIG. 8:**
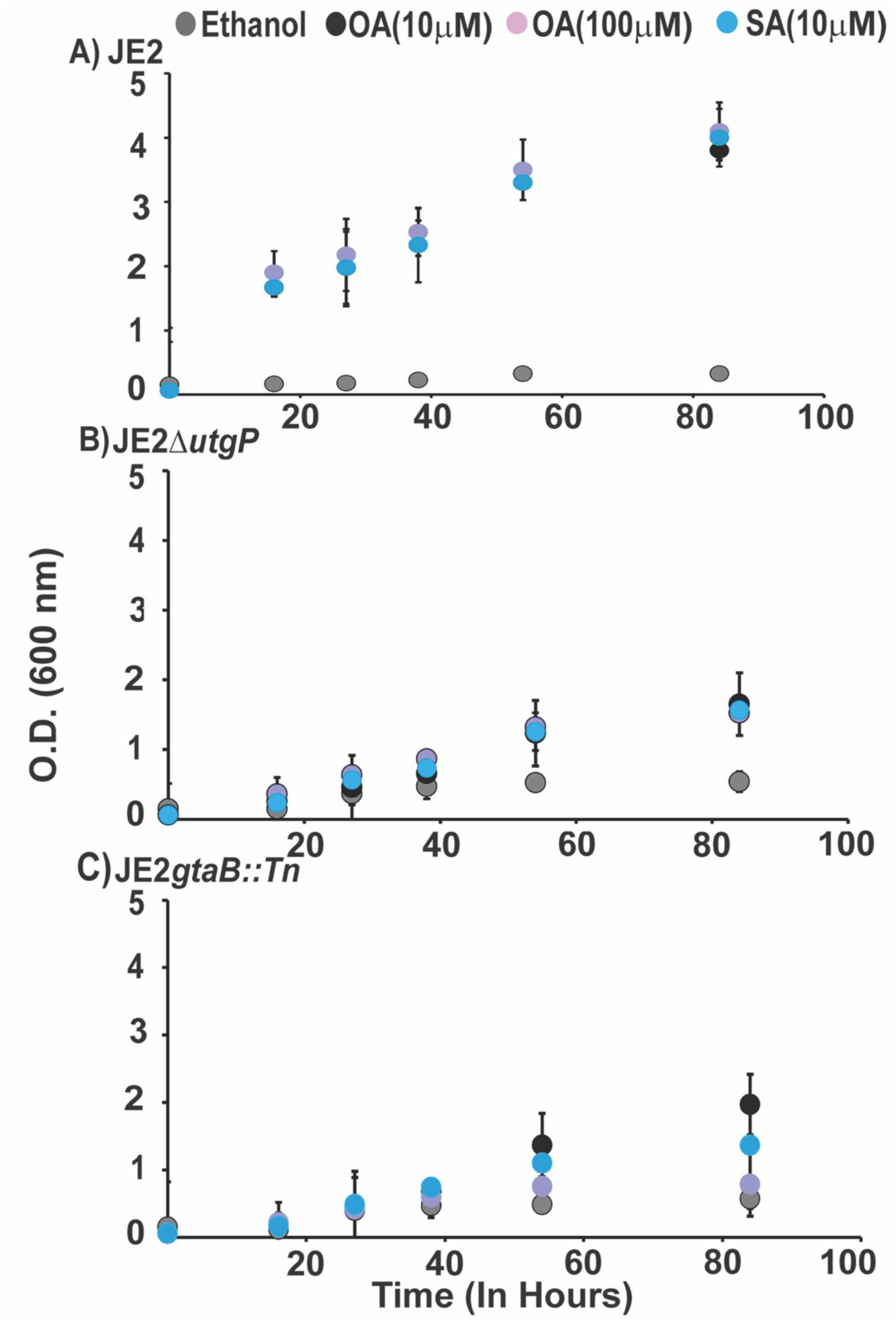
Glycolipid-deficient mutants exhibit reduced SCUFA-mediated growth stimulation at 12°C. Growth of **(A)** JE2 and the glycolipid-deficient mutants **(B)** JE2Δ*ugtP* and **(C)** JE2 *gtaB::Tn* in the presence of oleic acid (OA) at 10 (black circles) and 100 μM (lavender circles) and sapienic acid at 10 µM (SA, blue circles) at 12°C. Compared with the parent strain, both mutants showed markedly reduced growth stimulation by SCUFAs, indicating that glycolipids are important for membrane adaptation and growth at low temperature.

**FIG. 9:**
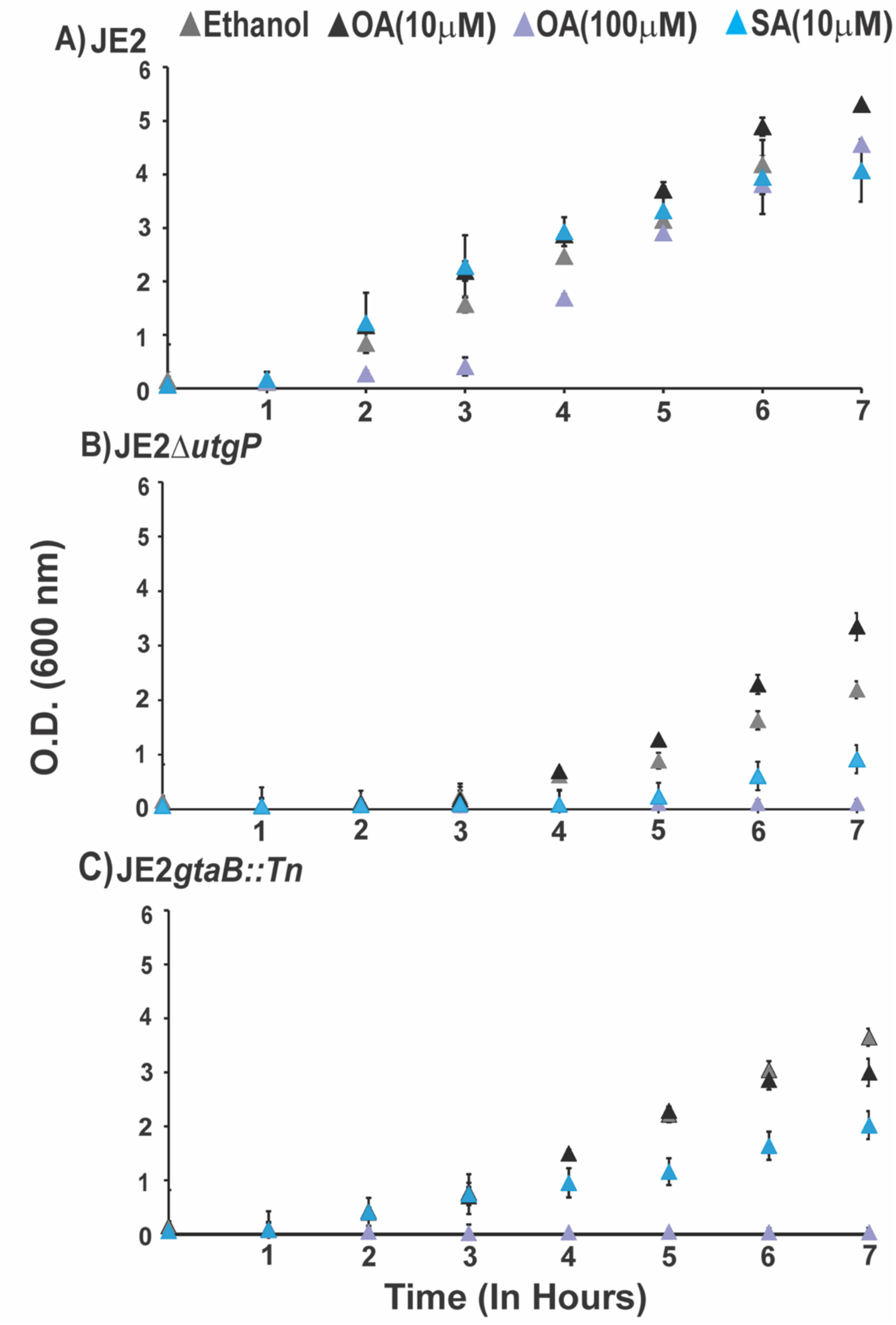
Glycolipid deficiency increases susceptibility to antimicrobial fatty acids at 37°C. Growth of **(A)** JE2 and the glycolipid-deficient mutants **(B)** JE2Δ*ugtP* and **(C)** JE2 *gtaB::Tn* in the presence of oleic acid (OA) at 10 (black triangles) and 100 µM (lavender triangles) and sapienic acid (SA, blue triangles) at 37 °C. While JE2 showed minimal inhibition, both glycolipid-deficient mutants were completely inhibited by 100 µM oleic acid and exhibited increased sensitivity to 10 µM sapienic acid.

## DISCUSSION

Our study revealed that the presence of antimicrobial SCUFAs in *S. aureus* cultures grown at low temperatures resulted in increased DGDG production and their preferential incorporation into the glycolipid DGDG, strong promotion of susceptilow-temperature growth and enhanced the production of the carotenoid staphyloxanthin. Lipidomics studies revealed that SCUFA incorporation into *S. aureus* membrane lipids was a feature of exposure of the organism to these molecules at concentrations below their MICs. Growth was promoted by the SCUFAs at 12°C, and this was dependent on their incorporation into membrane phospho- and glyco-lipids, because no growth stimulation occurred in a FakA mutant. Experiments with glycolipid-deficient mutants revealed that these glycolipids were important for low-temperature adaptation and tolerance of SCUFAs at higher temperatures.

### Preferential incorporation of SCUFAs into DGDGs over PGs at 12°C

At 12°C we observed that SCUFAs were preferentially enriched into the DGDG glycolipids. Previous analyses of DGDG fatty acid composition indicated that SCFAs, (C16:0, C18:0, C20:0) are preferentially linked to position *sn*-1 and *iso*-and *anteiso* C15:0 to the *sn*-2 position in cells grown in conventional laboratory media [23]. Although glycolipids are major lipid components of many Gram-positive bacteria, they have tended to be neglected in consideration of bacterial membrane composition and function. However recent papers recognize the role of glycolipids in streptococcal diabetic wound infection [24] and in the structure and function of the cytoplasmic membrane of *Clostridioides difficile* [25].

DGDGs are synthesized by the consecutive addition of two glucose units onto the *sn*-3 position of a DG by the enzyme YpfP, also known as UgtP **(FIG.1)**. DGDG serves a dual function in *S. aureus* as both a membrane lipid and a membrane anchor for lipoteichoic acid (LTA) [23] and the main source of DG in *S. aureus* derives from the synthesis of LTA, upon which LtaS transfers the glycerophosphate headgroup of a PG lipid to elongate the poly-glycerophosphate chain of LTA. The resulting DG, which is lethal if allowed to accumulate [26], can be used by either UgtP as a substrate for DGDG synthesis or can be phosphorylated by the diacylglycerol kinase DgkB [26, 27] to form phosphatidic acid (PA). The activity of DgkB is essential to the recycling of DG back into major phospholipid of *S. aureus* membranes, PG. However, there are alternate routes for the synthesis of PGs through the PlsX/Y/C pathway into which exogeneous FAs are directed **(FIG.1)**. DGDG is often quoted as representing 8 mol% of the total lipids of *S. aureus* [28, 29]. However, this is an underestimation in that LTA (6 mol%) and diacylglycerol (DG) (24 mol%) were included in the analysis, and these are not usually reported in the total lipid composition. Thus, quantitatively DGDG is a major *S. aureus* membrane lipid component. The enrichment of DGDGs with exogenous SCUFAs could therefore suggest that SCUFA-containing PGs are preferred substrates for UtgP and LtaS in contrast to the resulting DG remnants for recycling by DgkB. It is plausible that the only route available to transform a lethal accumulation of SCUFA-containing DG from LTA synthesis is through its conversion into DGDG.

### A significant role for DGDGs in low temperature adaptation

The presence of SCUFAs in the growth medium increased the total amount of DGDGs (saturated and unsaturated) in bacteria grown at 12°C. As noted above, the most affected endogenous DGDGs were those with acyl tail compositions of 15:0/15:0, 16:0/15:0, and 17:0/15:0 but the presence of SCUFAs in the 12°C growth conditions also restored the abundance of longer-tailed endogenous DGDGs, such as DGDG 33:0, 34:0, and 35:0, towards the levels of the ethanol control at 37°C. This SCUFA-driven stimulation of endogenous DGDGs was more significant than the direct incorporation of SCUFAs into DGDGs or PGs, which represented just 1/10^th^ to 1/100^th^ the abundance of the endogenous species, respectively. These data suggest that it is not solely the presence of SCUFA-containing lipids that stimulate the growth of *S. aureus* at low-temperatures, but perhaps their impact on the synthesis of specific lipid structures, DGDGs, instead. This is further supported by our findings that two glycolipid-deficient mutants showed a much-diminished growth response to SCUFAs at low temperatures. Our findings suggest that DGDG plays a major role in the adaptation of *S. aureus* to low temperatures. However, Δ*ugtP* deletion mutants are pleiotropic, lacking the DGDG anchor of LTA, which is replaced with PG, and the lipoteichoic acid produced is abnormally long [30, 31]. While at this point we are not aware of much evidence for glycolipids playing roles in bacterial low temperature adaptation, increasing the proportion of digalactosyl diglyceride has been associated with low temperature growth in the green algae *Dunaliella salina* [32], and this was proposed to increase membrane fluidity. However, in the psychrotrophic bacterium *Listeria monocytogenes,* a cold-sensitive mutant was found to have a transposon insertion in gene *lmo1078* encoding a UDP glucose pyrophosphorylase (GtaB). In Gram-positive bacteria, this enzyme catalyzes the formation of UDP glucose, the main purpose of which is to be a substrate for glycosylation of wall teichoic acid and the formation of DGDGs [33]. Interestingly, *L. monocytogenes* produced two structural variant forms of lipoteichoic acid, LTA1 and LTA2, with LTA1 predominating in cells grown at room temperature [34]. Furthermore, Jones et al. [35] showed that a glycolipid was only produced in *L. monocytogenes* growing at low temperatures (10°C).

The biophysical properties of glycolipids differ markedly from those of phospholipids. Long-chain glycolipids, in particular, can form inverted hexagonal or cubic phases. When mixed with bilayer-forming lipids, they increase the lateral pressure in the membrane center, which is attributed to the weak hydration and extensive hydrogen bonding of the glycosyl head groups [15, 36–38]. We previously found that supplementation of *S. aureus* with oleic acid led to the appearance of long-chain saturated and unsaturated glycolipids. Lipid extracts from supplemented cultures showed a high-temperature transition in differential scanning calorimetry, consistent with the formation of an inverted lipid phase that stabilizes the cell membrane [15, 39].

An interesting observation was made when the glycolipid-deficient mutants grown at 37°C were shown to be completely susceptible to 100 μM oleic acid, whereas the MIC for wild type *S. aureus* is > 500µM. It may be that incorporation of oleic acid into PG, CL, and LPG renders the membrane hyperfluid in the absence of DGDG. DGDG may be required in order to incorporate some of the SCUFAs and impart stability to the membrane.There is growing evidence for the role of DGDG in membrane resilience and survival of toxic molecules. *ugtP* is required for β-lactam resistance in methicillin-resistant *S. aureus* [40]. Multi-strain Tn-seq revealed that *ugtP* was an important determinant of tolerance of daptomycin exposure [41].

### Low temperature adaptation of *S. aureus* involves multiple strategies

When Sinensky [42] developed his concept of homeoviscous adaptation to growth temperature in *Escherichia coli*, he focused on the role of change in fatty acid composition to provide a constant viscosity at the growth temperature. Fatty acid shortening and increased biosynthesis of SCUFAs occurred as growth temperature dropped. Amongst the Bacillales order of the Firmicutes bacteria such as *Bacillus*, *Listeria*, *Staphylococcus* genus members, the fatty acids are a mixture of BCFAs and SCFAs [43], and these bacteria do not generally biosynthesize SCUFAs. In the absence of SCUFAs the main modes of homeoviscous adaptation are in fatty acid shortening and branched-chain switching from *iso* to *anteiso* [44, 45]. Similar changes in fatty acid composition have been described in *S. aureus* [46–48]. However, we recently showed that oleic acid was incorporated in large amounts by *S. aureus* when present in the growth medium, and this promoted the growth of the organism at low temperatures [14]. In the current study, we turned our attention to other SCUFAs that have antimicrobial properties but nevertheless promote the growth of *S. aureus* at 12°C at low concentrations. Analysis of the membrane lipid composition of *S. aureus* following growth with SCUFAs at 12°C and 37°C revealed significant changes in both the overall abundance of lipid species as well as the structures of their acyl tails. In general, low temperature growth stimulated the production of lipid species with shorter acyl tails, such as PG 15:0/15:0 and DGDG 15:0/15:0, regardless of the presence of SCUFAs. However, the presence of SCUFAs significantly increased the total amount of DGDGs (saturated and unsaturated) in bacteria grown at 12°C. It was an unexpected finding that the *trans* isomers of oleic acid and vaccenic acid also stimulated low temperature growth at 12°C. Fully saturated C18:0 fatty acid did not stimulate low temperature growth.

In addition to fatty acid shortening, branching switching, incorporation of SCUFAs, and enhanced biosynthesis of glycolipids, the combination of low temperature and SCUFAs is a potent inducer of enhanced staphyloxanthin production. Kenny et al. [48] reported that the staphyloxanthin biosynthesis operon was upregulated by challenge with linoleic acid. Joyce et al. [46] also showed increased staphyloxanthin production at 25°C, and Seel et al. [49] have shown increased staphyloxanthin production in *Staphylococcus xylosus* at 10°C. A carotenoid-deficient *S. aureus* mutant showed significantly compromised growth stimulation by SCUFAs at low temperature, suggesting that increased staphyloxanthin production may contribute to low temperature growth. Recent studies on the role of staphyloxanthin in membrane organization by Múnera-Jaramillo et al. [50] offer some insights into how staphyloxanthin may play a role in membrane adaptation to low temperatures. Due to the greater length of staphyloxanthin hydrocarbon chains compared to those of phospholipids or glycolipids, it is proposed that staphyloxanthin, which spans the bilayer, orients itself at different angles with respect to the phospholipids and disrupts the ordered packaging of the phospholipids in the gel phase, decreasing the transition temperature. It is proposed that phospholipid acyl chains retain disorder in the liquid crystalline phase in the presence of staphyloxanthin that crisscrosses the bilayer at angled orientations through the core of the membrane bilayer.

### Implication of promotion of low temperature growth by SCUFAs in foodborne *S. aureus* intoxications

*S. aureus* remains a major foodborne pathogen worldwide. Maintaining the integrity of the cold chain is essential to prevent the growth of the organism in the food matrix [52]. It has been suggested that enhanced growth of *S. aureus* via oleic acid at low temperatures could have implications for the growth of the organisms at food preservation temperatures [14, 15]. Enhancement of low temperature growth by exogenous SCUFAs has been demonstrated in the related Gram-positive microorganism *L. monocytogenes.* Touche et al. [52] have shown that C18:1Δ9 was incorporated into *L. monocytogenes* lipids when it was supplied in the medium in the form of Tween 80 (polysorbate 80), and this enhanced the growth of the organism at refrigeration temperatures. Flegler et al. [53] showed that in addition to the enhancement of low temperature growth by oleic acid (C18:1Δ9), linoleic acid (C18:2Δ9,12) and linolenic acid (C18:3Δ9,12,15) also enhanced growth at low temperatures. SCUFAs are present in low-fat food matrices at only a few microorganisms per gram of food, whereas high-fat animal fats and plant-derived oils may contain SCUFAs at grams per kilogram of total fat levels. Reformulation of food with mono- and polyunsaturated fats [55] for health benefits may inadvertently compromise refrigeration as a control of bacterial growth due to the incorporation of SCUFAs into their lipids.

### Implications of the studies for the mode of action of antimicrobial SCUFAs

The mechanisms of action of antimicrobial SCUFAs in *S. aureus* and other bacteria are incompletely understood [56], and may include a variety of effects on membrane and membrane-related processes. SCUFAs such as sapienic acid are thought to insert into the membrane and increase fluidity [57]. Accumulation of SCUFAs in the membrane can lead to the creation of pores and detergent-like solubilization of the membrane [58]. Palmitoleic acid treatment of *S. aureus* led to dye-binding to DNA, and release of ATP and proteins, indicating disruption of the permeability barrier function of the membrane [20]. Further effects of SCUFAs include disruption of the electron transport chain, uncoupling of oxidative phosphorylation, interference with fatty acid biosynthesis and lipid peroxidation, and auto-oxidation. We have shown that all the SCUFAs we have studied were incorporated into phospholipid and glycolipid structures. These SCUFA-containing lipids are likely to increase the fluidity of the membrane in their vicinity, and create what are known as regions of increased fluidity [59]. These regions of increased fluidity may facilitate the insertion of the free form of SCUFAs into the membrane. Support for this idea is the finding that inactivation of the FakAB pathway for the incorporation of exogenous fatty acids leads to increased resistance to SCUFAs [60]. Also compatible with this notion are the findings in the study of two synthetic antimicrobial SCUFAs, 2-hexadecynonic acid (2-HDA) and (*Z*)-2-allyl-3-bromo-2-hexadecenoic acid (DAT-51) [61]. 2-HDA primarily disrupted membranes by insertion and pore formation, whereas DAT-51 had more moderate membrane perturbation but also inhibited peptidoglycan biosynthesis. Interestingly, lipidomics revealed the incorporation of 2-HDA into *S. aureus* lipids, but not DAT-51. Sidders et al. [61] have shown that the antimicrobial SCUFA palmitoleic acid synergizes with vancomycin and potentiates its antibacterial activity in a wide variety of Gram-positive bacteria. Incorporation of palmitoleic acid into phospholipids and glycolipids may enhance the generation of membrane regions of increased fluidity containing membrane-bound cell wall biosynthetic intermediates, and loss of membrane integrity.

### Summary

In summary, all antimicrobial SCUFAs tested were incorporated into glycolipids and phospholipids when they were presented at sub-MICs in the growth medium. The SCUFAs were preferentially incorporated into DGDG over PG at 12°C. Growth at 12°C was stimulated by SCUFAs and accompanied by enhanced DGDG production. Glycolipid-deficient mutants showed a much-diminished growth response to SCUFAs at 12°C. The *S. aureus* lipidome is complex being composed of BCFAs, SCFAs, SCUFAs, phospholipids, glycolipids, staphyloxanthin, and LTA. The membrane lipid remodeling required to meet the stress of low temperature growth is more complex than the conventional fatty acyl-centric view of homeoviscous adaptation, and involves changes in fatty acid unsaturation, fatty acid shortening, and branching switching, as well as increased glycolipid and carotenoid production. It is expected that significant changes in the lipidome are necessary in responding to other environmental stresses including that of the host environment.

## EXPERIMENTAL PROCEDURES

### Bacterial strains used and growth conditions

The strains used in this investigation are shown in **SUPPLEMENTARY Table S2**. Unless indicated otherwise, the strains were grown overnight at 37°C in Tryptic Soy Broth (TSB) (BD Bacto, Fisher Scientific) with shaking. This culture was used to inoculate 96 well plates (Cell Pro) containing TSB supplemented with various fatty acids, triglycerides and cholesteryl esters to a starting OD_600_ of 0.05. The plates were incubated with shaking at 200 rpm at 12°C for the indicated times and growth was measured at an interval of 10 to 12 hours in the Tecan Infinite 200 pro plate reader. Alternatively, cultures of 50 ml were grown in 250 ml Erlenmeyer flasks with shaking at 200 rpm at the indicated temperatures.

### Lipids

Fatty acids, triglycerides, and cholesteryl esters were purchased from Larodan Research Grade Lipids or Cayman Chemicals. Stock solutions were prepared in 95% (v/v) ethanol and were stored at -20°C.

### Estimation of carotenoid content

Washed harvested cells were extracted with warm methanol (55°C) for 5 minutes and the OD_465_ of the supernatant was measured as described previously [14].

### Lipid Extraction

Lipid extraction was performed using a modified Bligh-Dyer extraction [63]. Bacterial pellets were washed with 2 mL of sterile water twice. During second washing, bacterial aliquots were taken for optical density at 600 nm. The remaining bacteria were pelleted and resuspended in 0.5 mL of HPLC grade water, transferred to a 5 mL glass extraction tube and sonicated for 30 minutes in an ice bath. Following sonication, 2 mL of chilled extraction solvent, 1:2 chloroform/methanol (*v/v*), was added to samples and vortexed for 5 minutes intermittently. Phase separation was achieved by adding an additional 0.5 mL of chloroform and 0.5 mL of water with 1 minute of vortexing prior to being centrifuged at 3,000 *x g* at 4C for 10 minutes. The bottom layer was isolated into a fresh extraction tube and vacuum-dried (Savant, ThermoScientific). Dried extracts were reconstituted in 0.5 mL of 1:1 chloroform/methanol and stored at -80°C until analysis.

### Liquid-Chromatography Mass Spectrometry Data Collection and Analysis

*Lipidomics* was performed using a Waters Acquity Ultra Performance Liquid Chromatography system coupled with Waters Synapt XS traveling-wave ion mobility-mass spectrometer. Lipid extracts were dried under vacuum and reconstituted in mobile phase A (MPA) with 500- or 25-fold dilutions for negative or positive mode analysis, respectively. Samples were maintained at 6°C throughout data collection, and 5 μL was injected for analysis. Chromatographic separation was performed utilizing a Waters Cortec hydrophilic interaction liquid chromatography column (2.1 x 100 mm, 1.6 mm). Separations were performed at 40°C with a flow rate of 0.5 mL/min. MPA was comprised of 95% acetonitrile and 5% water with 10 mM ammonium acetate. Mobile phase B (MPB) was comprised of 50% acetonitrile and 50% water with 10 mM ammonium acetate. The gradient was as follows: 0-0.5 min at 100% MPB, 0.5-5 min from 100% to 60% MPB, 5-5.5 min at 60% MPB, 5.5-6 min from 60% to 100% MPB, 6-7 min to equilibrate at 100% MPB.

Following chromatography, samples were ionized using electrospray ionization using the following settings: capillary voltage, ±2 kV; source temperature, 150 °C; desolvation temperature, 500 °C; desolvation gas flow, 1000 L/h; cone gas flow, 50 L/h. Ion mobility separation was performed in nitrogen (90 mL/min flow) with traveling wave settings of 550 m/s and 40V. Data was collected over 50-1,200 m/z with a 0.5 s scan time for full-scan MS1, data-independent MS/MS, and leucine enkephalin lock-mass functions. Targeted MS/MS acquisition was collected in a separate analysis to determine fatty acid tail composition (Quadrupole isolation window: 14.0; collision energy, ramped 40-60 V). Waters .raw files were imported into Progenesis QI software (v3.0, Waters/Nonlinear Dynamics). Data was lock-mass corrected and aligned to a pooled quality control sample prepared from equal parts of all samples. Peak picking was performed using default parameters, and data was normalized to total ion count. PGs [M-H]^-^ were evaluated from the negative mode data, while DGDGs [M+NH_4_]^+^ and LysylPGs [M+H]^+^ were evaluated from the positive mode data. All lipids were identified by accurate mass (<10 ppm tolerance) using a modified version of LipidPioneer [64] as well as agreement with HILIC retention times and MS/MS fragmentation patterns of standards [65, 66].

## Data availability

All mass spectrometry data is available from the MassIVE repository under accession number MSV000100160 (https://doi.org/doi:10.25345/C5V11W036). Reviewers may access the dataset with the password 12I5F3QT2A5cb87h.

## ACKNOWLEDGEMENTS

This work was supported by National Institutes of Health grants R01AI173144 to K.M.H., V.K.S., and B.J.W., and 1R03AI174033-01A1 to J.-U.D. S.P. was supported by Mockford-Thompson fellowships in 2024 and 2025. We thank Dr. Francis Alonzo (University of Illinois Chicago) for providing the *geh* deletion mutant.

## References

1. Vestergaard M, Frees D, Ingmer H (2019) Antibiotic Resistance and the MRSA Problem. Microbiology Spectrum 7:. 10.1128/microbiolspec.gpp3-0057-2018

2. Hennekinne JA, De Buyser ML, Dragacci S (2012) Staphylococcus aureus and its food poisoning toxins: Characterization and outbreak investigation. FEMS Microbiology Reviews 36:. 10.1111/j.1574-6976.2011.00311.x

3. J Y, Co R (2017) Exogenous fatty acid metabolism in bacteria. Biochimie 141:. 10.1016/j.biochi.2017.06.015

4. Brinster S, Lamberet G, Staels B, et al (2009) Type II fatty acid synthesis is not a suitable antibiotic target for Gram-positive pathogens. Nature 458:83–86. 10.1038/nature07772

5. Kénanian G, Morvan C, Weckel A, et al (2019) Permissive Fatty Acid Incorporation Promotes Staphylococcal Adaptation to FASII Antibiotics in Host Environments. Cell Reports 29:3974–3982.e4. 10.1016/j.celrep.2019.11.071

6. Waters JK, Eijkelkamp BA (2024) Bacterial acquisition of host fatty acids has far-reaching implications on virulence. Microbiology and Molecular Biology Reviews 88:e00126–24

7. Parsons JB, Frank MW, Subramanian C, et al (2011) Metabolic basis for the differential susceptibility of Gram-positive pathogens to fatty acid synthesis inhibitors. Proc Natl Acad Sci U S A 108:15378–15383. 10.1073/pnas.1109208108

8. Frank MW, Yao J, Batte JL, et al (2020) Host fatty acid utilization by staphylococcus aureus at the infection site. mBio 11:. 10.1128/mBio.00920-20

9. Parsons JB, Broussard TC, Bose JL, et al (2014) Identification of a two-component fatty acid kinase responsible for host fatty acid incorporation by Staphylococcus aureus. Proceedings of the National Academy of Sciences of the United States of America 111:. 10.1073/pnas.1408797111

10. Hines KM, Alvarado G, Chen X, et al (2020) Lipidomic and Ultrastructural Characterization of the Cell Envelope of Staphylococcus aureus Grown in the Presence of Human Serum. mSphere 5:. 10.1128/msphere.00339-20

11. Pruitt EL, Zhang R, Ross DH, et al (2023) Elucidating the impact of bacterial lipases, human serum albumin, and FASII inhibition on the utilization of exogenous fatty acids by Staphylococcus aureus. mSphere 8:. 10.1128/msphere.00368-23

12. Sen S, Sirobhushanam S, Johnson SR, et al (2016) Growth-environment dependent modulation of Staphylococcus aureus branched-chain to straight-chain fatty acid ratio and incorporation of unsaturated fatty acids. PLoS ONE 11:. 10.1371/journal.pone.0165300

13. Delekta PC, Shook JC, Lydic TA, et al (2018) Staphylococcus aureus utilizes host-derived lipoprotein particles as sources of fatty acids. Journal of Bacteriology 200:. 10.1128/JB.00728-17

14. Barbarek SC, Shah R, Paul S, et al (2024) Lipidomics of homeoviscous adaptation to low temperatures in Staphylococcus aureus utilizing exogenous straight-chain unsaturated fatty acids. Journal of Bacteriology 206:e00187–24. 10.1128/jb.00187-24

15. Raskovic D, Alvarado G, Hines KM, et al (2025) Growth of Staphylococcus aureus in the presence of oleic acid shifts the glycolipid fatty acid profile and increases resistance to antimicrobial peptides. Biochim Biophys Acta Biomembr 1867:184395. 10.1016/j.bbamem.2024.184395

16. Fischer CL (2020) Antimicrobial Activity of Host-Derived Lipids. Antibiotics 9:75. 10.3390/antibiotics9020075

17. Mayo Clinic Laboratories, Endocrinology Catalog (Test ID: PFAPC) (2026) Fatty Acid Profile, Comprehensive (C8-C26), Plasma. https://endocrinology.testcatalog.org/show/PFAPE

18. Leekumjorn S, Cho HJ, Wu Y, et al (2009) The Role of Fatty Acid Unsaturation in Minimizing Biophysical Changes on the Structure and Local Effects of Bilayer Membranes. Biochim Biophys Acta 1788:1508–1516. 10.1016/j.bbamem.2009.04.002

19. Schmitt M, Schuler-Schmid U, Schmidt-Lorenz W (1990) Temperature limits of growth, TNase and enterotoxin production of Staphylococcus aureus strains isolated from foods. Int J Food Microbiol 11:1–19. 10.1016/0168-1605(90)90036-5

20. Parsons JB, Yao J, Frank MW, et al (2012) Membrane disruption by antimicrobial fatty acids releases low-molecular-weight proteins from staphylococcus aureus. Journal of Bacteriology 194:. 10.1128/JB.00743-12

21. Cadieux B, Vijayakumaran V, Bernards MA, et al (2014) Role of lipase from community-associated methicillin-resistant Staphylococcus aureus strain USA300 in hydrolyzing triglycerides into growth-inhibitory free fatty acids. J Bacteriol 196:4044–4056. 10.1128/JB.02044-14

22. Braungardt H, Singh VK (2019) Impact of Deficiencies in Branched-Chain Fatty Acids and Staphyloxanthin in Staphylococcus aureus. Biomed Res Int 2019:2603435. 10.1155/2019/2603435

23. Fischer W (1994) Lipoteichoic acid and lipids in the membrane of Staphylococcus aureus. Med Microbiol Immunol 183:61–76. 10.1007/BF00277157

24. Joyce LR, Akbari MS, Nguyen DT, et al (2025) Global genome analysis identifies glycolipids and lipoteichoic acid alanylation as contributors to group B streptococcal diabetic wound infection. Cell Rep 44:116510. 10.1016/j.celrep.2025.116510

25. Zbylicki BR, Cochran S, Weiss DS, Ellermeier CD (2025) Identification of two glycosyltransferases required for synthesis of membrane glycolipids in Clostridioides difficile. mBio 16:e03512–24. 10.1128/mbio.03512-24

26. Miller DJ, Jerga A, Rock CO, White SW (2008) Analysis of the Staphylococcus aureus DgkB Structure Reveals a Common Catalytic Mechanism for the Soluble Diacylglycerol Kinases. Structure 16:1036–1046. 10.1016/j.str.2008.03.019

27. Mychack A, Evans D, Gilles T, et al (2025) Staphylococcus aureus uses a GGDEF protein to recruit diacylglycerol kinase to the membrane for lipid recycling. Proceedings of the National Academy of Sciences 122:e2414696122. 10.1073/pnas.2414696122

28. Koch HU, Haas R, Fischer W (1984) The role of lipoteichoic acid biosynthesis in membrane lipid metabolism of growing Staphylococcus aureus. European Journal of Biochemistry 138:357–363. 10.1111/j.1432-1033.1984.tb07923.x

29. Kiriukhin MY, Debabov DV, Shinabarger DL, Neuhaus FC (2001) Biosynthesis of the Glycolipid Anchor in Lipoteichoic Acid of Staphylococcus aureus RN4220: Role of YpfP, the Diglucosyldiacylglycerol Synthase. Journal of Bacteriology 183:3506–3514. 10.1128/jb.183.11.3506-3514.2001

30. Hesser AR, Matano LM, Vickery CR, et al (2020) The Length of Lipoteichoic Acid Polymers Controls Staphylococcus aureus Cell Size and Envelope Integrity. Journal of Bacteriology 202:10.1128/jb.00149-20. 10.1128/jb.00149-20

31. Gründling A, Schneewind O (2007) Genes Required for Glycolipid Synthesis and Lipoteichoic Acid Anchoring in Staphylococcus aureus. Journal of Bacteriology 189:2521–2530. 10.1128/jb.01683-06

32. Lynch DV, Thompson Jr GA (1982) Low temperature-induced alterations in the chloroplast and microsomal membranes of Dunaliella salina. Plant physiology 69:1369–1375

33. Reichmann NT, Gründling A (2011) Location, synthesis and function of glycolipids and polyglycerolphosphate lipoteichoic acid in Gram-positive bacteria of the phylum Firmicutes. FEMS Microbiol Lett 319:97–105. 10.1111/j.1574-6968.2011.02260.x

34. Dehus O, Pfitzenmaier M, Stuebs G, et al (2011) Growth temperature-dependent expression of structural variants of Listeria monocytogenes lipoteichoic acid. Immunobiology 216:24–31. 10.1016/j.imbio.2010.03.008

35. Jones CE, Shama G, Jones D, et al (1997) Physiological and biochemical studies on psychrotolerance in Listeria monocytogenes. J Appl Microbiol 83:31–35. 10.1046/j.1365-2672.1997.d01-391.x

36. Hinz HJ, Kuttenreich H, Meyer R, et al (1991) Stereochemistry and size of sugar head groups determine structure and phase behavior of glycolipid membranes: densitometric, calorimetric, and X-ray studies. Biochemistry 30:5125–5138. 10.1021/bi00235a003

37. Mannock DA, Collins MD, Kreichbaum M, et al (2007) The thermotropic phase behaviour and phase structure of a homologous series of racemic beta-D-galactosyl dialkylglycerols studied by differential scanning calorimetry and X-ray diffraction. Chem Phys Lipids 148:26–50. 10.1016/j.chemphyslip.2007.04.004

38. Wizert A, Iskander DR, Cwiklik L (2017) Interaction of lysozyme with a tear film lipid layer model: A molecular dynamics simulation study. Biochim Biophys Acta Biomembr 1859:2289–2296. 10.1016/j.bbamem.2017.08.015

39. Osterberg F, Rilfors L, Wieslander A, et al (1995) Lipid extracts from membranes of Acholeplasma laidlawii A grown with different fatty acids have a nearly constant spontaneous curvature. Biochim Biophys Acta 1257:18–24. 10.1016/0005-2760(95)00042-b

40. Muscato JD, Morris HG, Mychack A, et al (2022) Rapid Inhibitor Discovery by Exploiting Synthetic Lethality. J Am Chem Soc 144:3696–3705. 10.1021/jacs.1c12697

41. Coe KA, Lee W, Stone MC, et al (2019) Multi-strain Tn-Seq reveals common daptomycin resistance determinants in Staphylococcus aureus. PLOS Pathogens 15:e1007862. 10.1371/journal.ppat.1007862

42. Sinensky M (1974) Homeoviscous Adaptation—A Homeostatic Process that Regulates the Viscosity of Membrane Lipids in Escherichia coli. Proceedings of the National Academy of Sciences 71:522–525. 10.1073/pnas.71.2.522

43. Kaneda T (1971) Factors affecting the relative ratio of fatty acids in Bacillus cereus. Can J Microbiol 17:269–275. 10.1139/m71-045

44. Annous BA, Becker LA, Bayles DO, et al (1997) Critical role of anteiso-C15:0 fatty acid in the growth of Listeria monocytogenes at low temperatures. Applied and Environmental Microbiology. 10.1128/aem.63.10.3887-3894.1997

45. Suutari M, Laakso S (1994) Microbial fatty acids and thermal adaptation. Crit Rev Microbiol 20:285–328. 10.3109/10408419409113560

46. Wang L-H, Wang M-S, Zeng X-A, Liu Z-W (2016) Temperature-mediated variations in cellular membrane fatty acid composition of Staphylococcus aureus in resistance to pulsed electric fields. Biochim Biophys Acta 1858:1791–1800. 10.1016/j.bbamem.2016.05.003

47. Joyce GH, Hammond RK, White DC (1970) Changes in membrane lipid composition in exponentially growing Staphylococcus aureus during the shift from 37 to 25 C. J Bacteriol 104:323–330. 10.1128/jb.104.1.323-330.1970

48. Singh VK, Hattangady DS, Giotis ES, et al (2008) Insertional inactivation of branched-chain α-keto acid dehydrogenase in Staphylococcus aureus leads to decreased branched-chain membrane fatty acid content and increased susceptibility to certain stresses. Applied and Environmental Microbiology 74:. 10.1128/AEM.00882-08

49. Kenny JG, Ward D, Josefsson E, et al (2009) The Staphylococcus aureus Response to Unsaturated Long Chain Free Fatty Acids: Survival Mechanisms and Virulence Implications. PLOS ONE 4:e4344. 10.1371/journal.pone.0004344

50. Seel W, Baust D, Sons D, et al (2020) Carotenoids are used as regulators for membrane fluidity by Staphylococcus xylosus. Sci Rep 10:330. 10.1038/s41598-019-57006-5

51. Múnera-Jaramillo J, López G-D, Suesca E, et al (2024) The role of staphyloxanthin in the regulation of membrane biophysical properties in *Staphylococcus aureus*. Biochimica et Biophysica Acta (BBA) - Biomembranes 1866:184288. 10.1016/j.bbamem.2024.184288

52. Bennett SD, Sodha SV, Ayers TL, et al (2018) Produce-associated foodborne disease outbreaks, USA, 1998-2013. Epidemiol Infect 146:1397–1406. 10.1017/S0950268818001620

53. Touche C, Hamchaoui S, Quilleré A, et al (2023) Growth of Listeria monocytogenes is promoted at low temperature when exogenous unsaturated fatty acids are incorporated in its membrane. Food Microbiol 110:104170. 10.1016/j.fm.2022.104170

54. Flegler A, Iswara J, Mänz AT, et al (2022) Exogenous fatty acids affect membrane properties and cold adaptation of Listeria monocytogenes. Sci Rep 12:1499. 10.1038/s41598-022-05548-6

55. Onyeaka H, Nwaiwu O, Obileke K, et al (2023) Global nutritional challenges of reformulated food: A review. Food Sci Nutr 11:2483–2499. 10.1002/fsn3.3286

56. Douglas EJA, Palk N, Rudolph ER, Laabei M (2025) Anti-staphylococcal fatty acids: mode of action, bacterial resistance and implications for therapeutic application. Microbiology 171:001563. 10.1099/mic.0.001563

57. Cartron ML, England SR, Chiriac AI, et al (2014) Bactericidal Activity of the Human Skin Fatty Acid cis-6-Hexadecanoic Acid on Staphylococcus aureus. Antimicrob Agents Chemother 58:3599–3609. 10.1128/AAC.01043-13

58. Arouri A, Mouritsen OG (2013) Membrane-perturbing effect of fatty acids and lysolipids. Prog Lipid Res 52:130–140. 10.1016/j.plipres.2012.09.002

59. Müller A, Wenzel M, Strahl H, et al (2016) Daptomycin inhibits cell envelope synthesis by interfering with fluid membrane microdomains. Proceedings of the National Academy of Sciences 113:E7077–E7086. 10.1073/pnas.1611173113

60. Krute CN, Ridder MJ, Seawell NA, Bose JL (2019) Inactivation of the exogenous fatty acid utilization pathway leads to increased resistance to unsaturated fatty acids in Staphylococcus aureus. Microbiology (Reading) 165:197–207. 10.1099/mic.0.000757

61. Villarini-Torres J, Casillas-Vargas G, Shayeb K, et al (2025) Dual Mechanisms of Action of a Halogenated Allyl Fatty Acid against Methicillin-Resistant Staphylococcus aureus. ACS Omega 10:55769–55786. 10.1021/acsomega.5c07162

62. Sidders AE, Kedziora KM, Arts M, et al (2023) Antibiotic-induced accumulation of lipid II synergizes with antimicrobial fatty acids to eradicate bacterial populations. eLife 12:e80246. 10.7554/eLife.80246

63. Bligh EG, Dyer WJ (1959) A rapid method of total lipid extraction and purification. Can J Biochem Physiol 37:911–917. 10.1139/o59-099

64. Ulmer CZ, Koelmel JP, Ragland JM, et al (2017) LipidPioneer : A Comprehensive User-Generated Exact Mass Template for Lipidomics. J Am Soc Mass Spectrom 28:562–565. 10.1007/s13361-016-1579-6

65. Freeman C, Hynds HM, Carpenter JM, et al (2021) Revealing Fatty Acid Heterogeneity in Staphylococcal Lipids with Isotope Labeling and RPLC–IM–MS. J Am Soc Mass Spectrom 32:2376–2385. 10.1021/jasms.1c00092

66. Bimpeh K, Hines KM (2023) A rapid single-phase extraction for polar staphylococcal lipids. Anal Bioanal Chem 415:4591–4602. 10.1007/s00216-023-04758-9

