## Supplementary Figures for "Exogenous antimicrobial straight-chain unsaturated fatty acids promote low-temperature growth of *Staphylococcus aureus*"

### SUPPLEMENTARY MATERIALS

#### Exogenous antimicrobial straight-chain unsaturated fatty acids increase diglucosyldiacylglyceride and staphyloxanthin levels to promote low-temperature growth of Staphylococcus aureus.

Sharanya Paul[1], David Brewer[2], Matthew W. Frank[3], Arunachalam Muthaiyan[4], Vineet K. Singh[5], Antje Pokorny[6], Kelly M. Hines[2], Jan-Ulrik Dahl[1], and Brian J. Wilkinson[1],*

#### SUPPLEMENTARY RESULTS

**Impact of SCUFAs on PG and DGDG species at 37°C.** Notably, the incorporation of any monounsaturated SCUFAs and linoleic acid had no impact on the saturated PG population (**FIG. 6**). However, arachidonic acid was found to suppress lower chain length saturated PGs like PG 30:0 and PG 31:0. The incorporation of exogenous SCUFAs was seen through the occurrence of SCUFA-specific unsaturated PGs. Out of PG 31:1 (16:1/15:0), which was only produced in the presence of sapienic and palmitoleic acid, all monounsaturated SCUFAs produced a similar unsaturated PG profile dominated by PG 33:1 (18:1/15:0) followed by PG 35:1 (20:1/15:0) (**FIG. 6C**). As these are the dominant monounsaturated PGs, this points towards a potential preference for the incorporation of FA 18:1 over FA 16:1 species into the acyl tails of PGs. Trace levels of PG 34:2 (18:1/16:1) were found with palmitoleic acid from incorporation into both the *sn-1* and *sn-2* position. While other even-chain length mono-unsaturated PG species associated with a FA 14:0 at the *sn-2* position, these made up a fairly minor portion of the overall lipidome. Linoleic acid was found to produce PG 33:2 (18:2/15:0) and PG 35:2 (20:2/15:0) as its major unsaturated species (**FIG. 6C**). Unlike its mono-unsaturated counterparts, both FA 18:2 and its first elongation to FA 20:2 were incorporated equally into PGs. Incorporation into even chain-length PGs was found to be more prominent with linoleic acid in comparison to its counterparts. Arachidonic acid was only found to produce PG 35:4 (20:4/15:0).

In addition to impacting the PG population, supplementation with exogenous SCUFAs were found to affect the DGDGs population (**FIG. 7**). As with PGs, the incorporation of SCUFAs into the DGDG population was seen. Species such as DGDG 33:1 and 35:1 were seen with supplementation for all monounsaturated SCUFAs. Elevated levels of these species were seen with sapienic and palmitoleic acid over oleic acid, and DGDG 31:1 was seen preferentially with sapienic acid over palmitoleic acid. Double incorporation of palmitoleic acid was seen through the formation of DGDG 34:2 and 36:2. Incorporation of linoleic acid was shown though DGDG 33:2, DGDG 35:2, and DGDG 37:2 with trends mimicking their PG counterparts. Arachidonic acid was found to produce DGDG 35:4 and 37:4, a species not found within the PG population. Comparing the total intensity of DGDGs across all conditions showed no significant increases in saturated DGDGs. Higher levels of polyunsaturated DGDGs were also seen with palmitoleic acid over sapienic and oleic acid. Similar levels of polyunsaturated DGDGs were seen with arachidonic acid compared to sapienic and palmitoleic acid; however, linoleic acid showed a higher level of unsaturated DGDGs compared to all other conditions. Overall, the proportions of unsaturated DGDG species with the different fatty acids were similar to the proportions of unsaturated PG species **(SUPPLEMENTARY FIG. S3)**.

**Impact of SCUFAs on PG species at 12°C.** Low-temperature growth promoted a shift towards PGs species with shorter acyl tails, as demonstrated by the significant increase in PG 30:0 (likely PG 15:0/15:0) in all bacteria grown at 12°C (**FIG. 6B**). This change was accompanied by a decrease in PGs with long-chain acyl tails, such as PG 35:0 (likely 20:0/15:0) and PG 33:0 (likely 18:0/15:0). This trend was independent of the presence or absence of SCUFAs as both the ethanol control and SCUFA-supplemented conditions followed the same pattern. However, PG 32:0 (likely 17:0/15:0) was elevated only in the bacteria that had been provided with SCUFAs during low-temperature growth and no difference was observed in the ethanol control between 12°C and 37°C. The incorporation of exogenous SCUFAs into staphylococcal PGs was greater in the bacteria grown at 37°C regardless of the specific SCUFA provided, but the overall amounts of unsaturated PGs was nearly 100-fold lower than the endogenous saturated species. The largest difference in SCUFA incorporation between the two growth temperatures was observed with linoleic acid, where the abundance of PGs containing linoleate-derived acyl tails was 75- to 4-fold higher at 37°C than 12°C. While the incorporation of arachidonic acid into PGs was also detected, the difference between 12°C and 37°C was less than that of linoleic acid. The sole exception to the observation of elevated SCUFA uptake at 37°C was the ethanol control condition in which bacteria produced PG species containing SCUFAs at 12°C even in the absence of supplementation of the medium with exogenous SCUFAs. For PGs with shorter acyl tails, such as PG 31:1 and PG 32:1, the abundance in the 12°C ethanol control was equal to or slightly higher than the conditions in which SCUFAs were provided in the broth **(FIG. 6, SUPPLEMENTARY FIG. S3)**. These data suggest that *S. aureus* was able to scavenge SCUFAs from the medium to support its growth at 12°C. Although very low in abundance, there was also evidence that exogenous SCUFAs were esterified at both glycerol backbone positions, including PG 36:2 in the 12°C ethanol control and the formation of PG 34:2 from palmitoleic acid at 37°C **(SUPPLEMENTARY FIG. S3)**.

**Lipidomic impact of C18:1 Fatty Acids.** Lipidomics analysis was performed to identify the changes in lipid classes and individual lipids species driven by growth with oleic acid and its isomers. At 12°C, SCUFA supplementation enhanced the formation of saturated PGs with 32 carbons or more, with oleic and *cis*-vaccenic acid having the greatest impact **(SUPPLEMENTARY FIG. S5-7)**. Additionally, while the impacts of both *trans*-fatty acids were identical, oleic acid enhanced saturated PG levels more than its counterpart, *cis*-vaccenic acid. Conversely, elaidic and *trans*-vaccenic acid suppressed all saturated PGs at 37°C while oleic and *cis*-vaccenic acid had no effect on the major saturated PG species (*i.e.,* PG 32:0 to PG 35:0). As with their impact on saturated PGs, the 18:1 SCUFA isomers were incorporated into PG species to varying degrees between the two growth temperatures. Overall incorporation of the 18:1 SCUFAs was greater at 12°C compared to 37°C, with the *trans*-geometry isomers (elaidic and *trans*-vaccenic) having the highest incorporation into PGs **(SUPPLEMENTARY FIG. S7)**. However, the incorporation of the *trans*-geometry isomers was far lower at 37°C relative to both 12°C and the amount of incorporation for the cis-geometry isomers at 37°C. Based on the similarities in the profile of PG 35:1 at 12°C for all isomers, there was no significant difference in elongation of the SCUFAs from 18 to 20-carbons due to their double bond geometries. The incorporation of two SCUFAs into PGs was minimal compared to the amounts of monounsaturated PGs **(SUPPLEMENTARY FIG. S7)**, but the direct incorporation of two *cis*-vaccenic acid acyl tails at 12°C was more prevalent than any other combination of SCUFAs and growth temperatures.

**SUPPLEMENTARY FIGURES & TABLES**

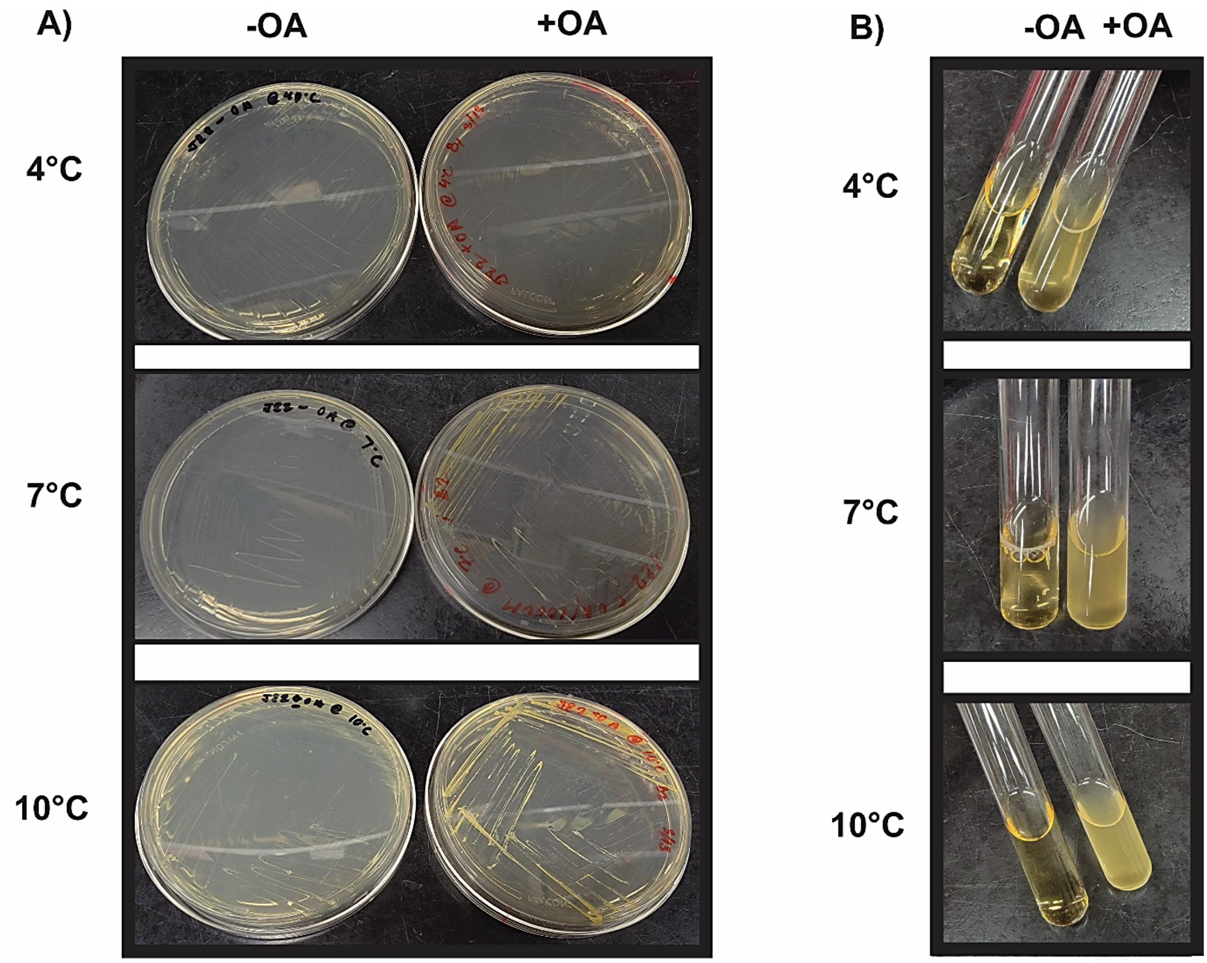


**SUPPLEMENTARY FIG S1. Supplementation of exogenous OA reduces the minimum growth temperature of JE2.** JE2 cultures were incubated in tryptic soy broth (TSB) at 10 °C, 7 °C, and 4 °C, with and without 100 μM OA supplementation. Growth was assessed on both **A)** solid and **B)** liquid media on day 30. Enhanced growth is evident in oleic acid-supplemented conditions compared to EtOH controls. One representative image of each condition shown.

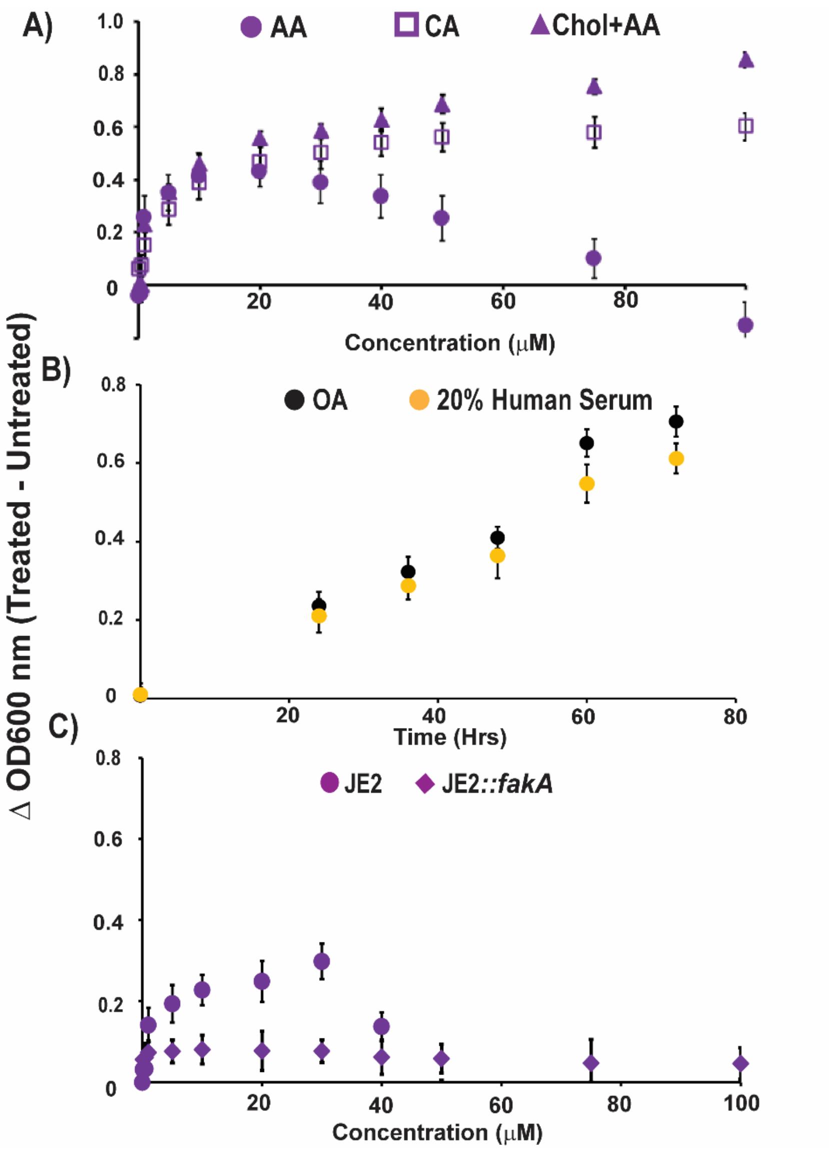


**SUPPLEMENTARY FIG S2.** Overnight cultures of JE2 were diluted into fresh TSB supplemented with increasing concentrations of **(A)** arachidonic acid (AA, filled circles), cholesteryl arachidonate (CA, squares) or co-incubation of arachidonic acid with 300 $\mu$M cholesterol (Chol + AA, filled triangles) to an $OD_{600}$ ~0.05 and cultivated for at 12˚C. $OD_{600}$ was determined after 5 days and normalized by subtracting the $OD_{600}$ of the ethanol-only control in case of AA and CA and for the co-incubation (Chol+AA) the $OD_{600}$ was determined after 5 days and normalized by subtracting the $OD_{600}$ of the cholesterol-only control, (n=3, ±S.D.). **(B)** TSB supplemented with either 20% human serum (yellow circles) or 100 µM oleic acid (black circles) was inoculated with JE2 and cultivated in 96-well plates at 12°C for 76 hrs. Growth was normalized by subtracting the $OD_{600}$ of the ethanol-only control, (n=2, ±S.D.). **(C)** JE2 (circles) and JE2 *fakA::Tn* (diamonds) were grown in TSB media in the presence and absence of the indicated concentrations of arachidonic acid. $OD_{600}$ was determined after 5 days and normalized by subtracting the $OD_{600}$ of the ethanol-only control, (n=3, ±S.D.).

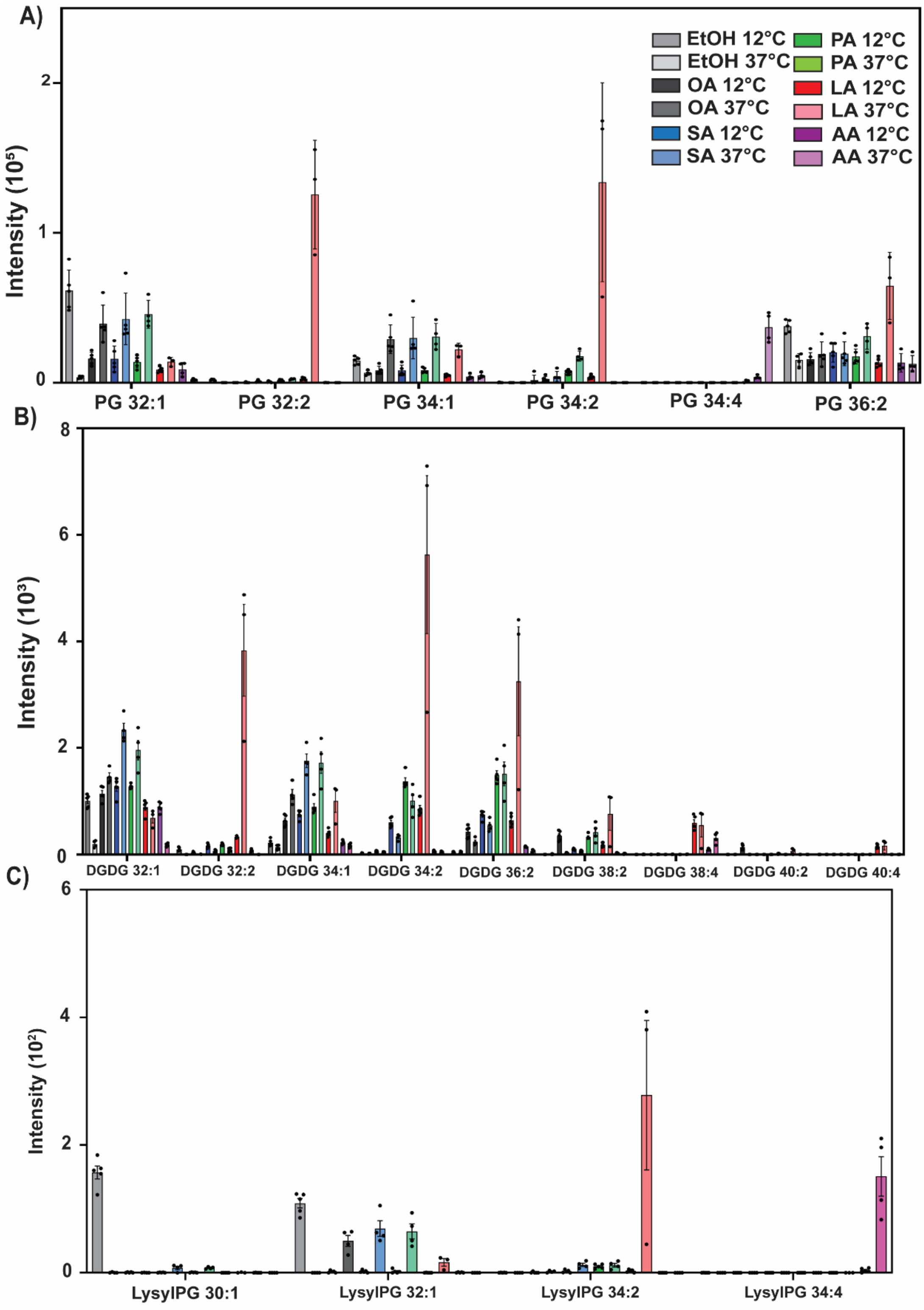


**SUPPLEMENTARY FIG S3. Effects of exogenous SCUFA supplementation on PG, DGDG, and Lysyl-PG lipids at 12°C and 37°C.** *S. aureus* strain JE2 was grown in the presence and absence 10 µM of the indicated SCUFAs at 12 °C and 37°C, respectively. **(A)** Impact on native unsaturated PG species. **(B)** Impact on unsaturated DGDG species. **(C)** Impact on unsaturated LysylPG species, (n=5, ±S.D.).

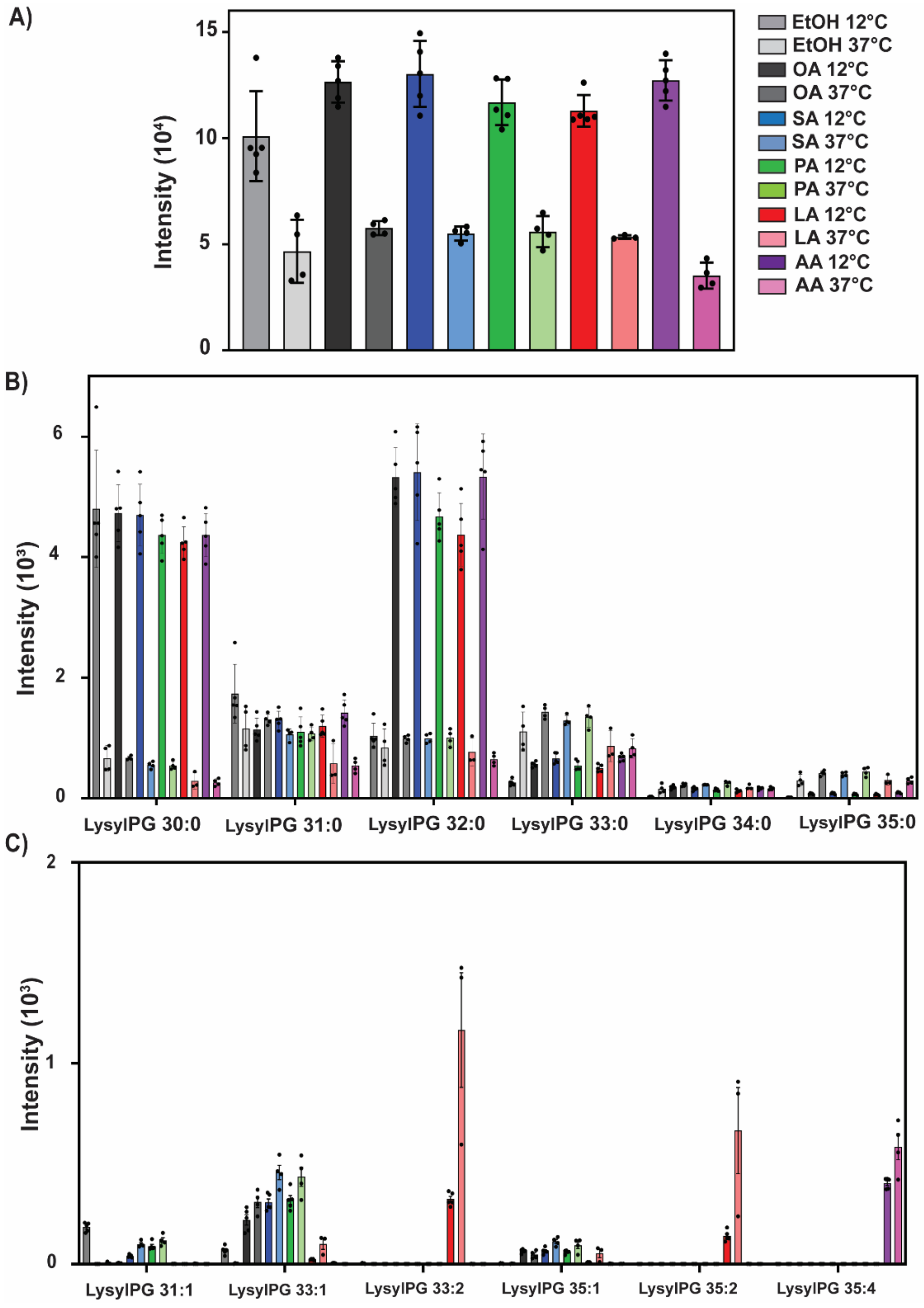


**SUPPLEMENTARY FIG S4. Effects of exogenous SCUFA supplementation on Lysyl-PG lipids at 12°C and 37°C.** *S. aureus* strain JE2 was grown in the presence and absence of 10 μM of the indicated SCUFAs at 12°C and 37°C, respectively. **(A)** Total Lysyl-PG abundance across all species. **(B)** Impact on native saturated Lysyl-PG species. **(C)** Impact on unsaturated Lysy-PG species, (n=5, ±S.D.).

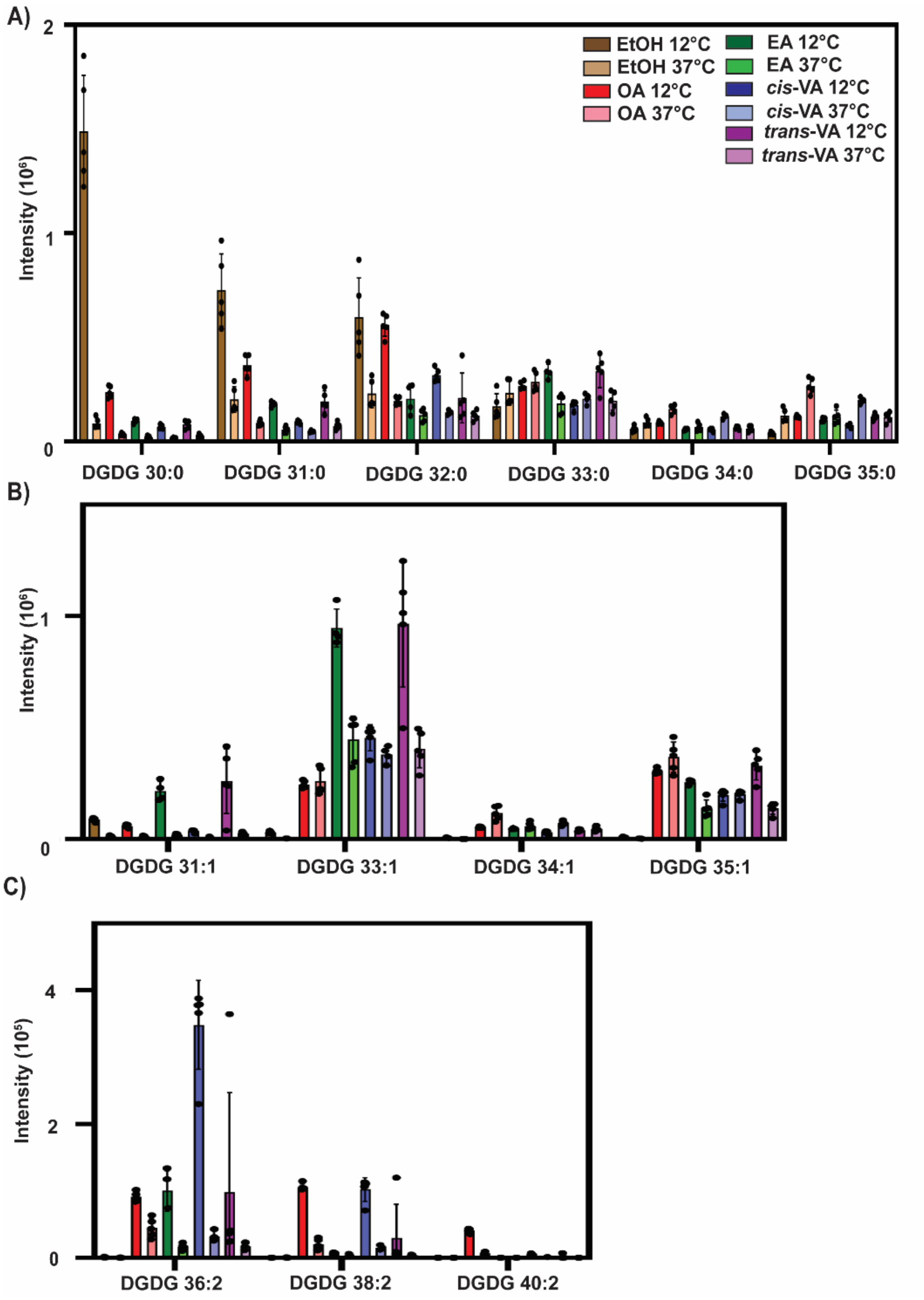


**Supplementary FIG S5:** Lipidomics analysis of DGDGs when supplemented with 100 μM of oleic acid (9Z), elaidic acid (9E), cis-vaccenic acid (11Z), or trans-vaccenic acid (11E) at 12 °C and 37°C. **(A)** DGDGs containing only saturated fatty acids. **(B)** High-abundance DGDGs containing one unsaturated fatty acid. **(C)** Low-abundance DGDGs containing two unsaturated fatty acids, (n=5, ±S.D.).

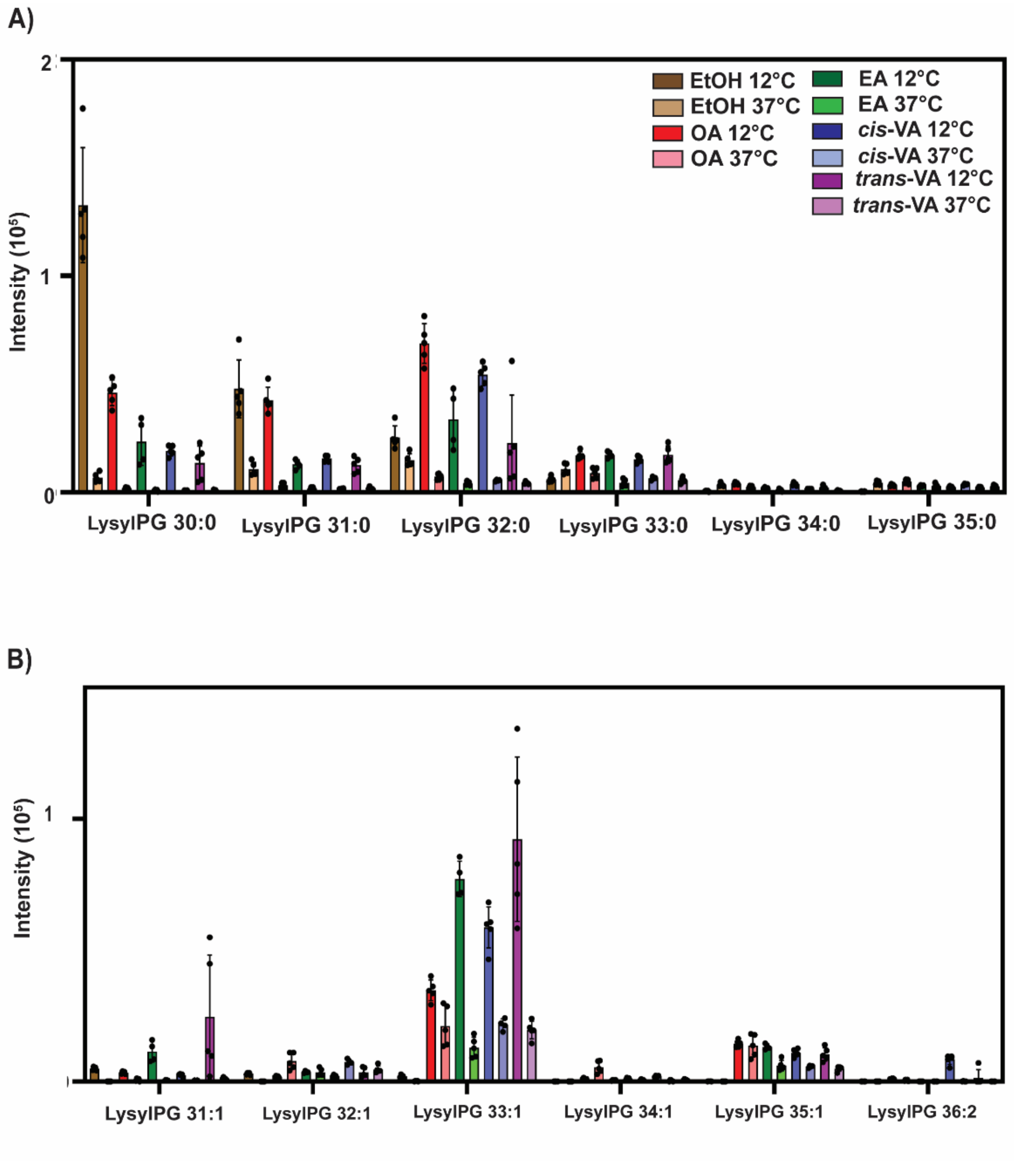


**SUPPLEMENTARY FIG S6. Effects of exogenous SCUFA supplementation on LysylPG lipids at 12°C and 37°C.** *S. aureus* strain JE2 was grown in the presence and absence 100 μM of oleic acid, elaidic acid, *cis*-vaccenic acid, or *trans*-vaccenic acid at 12 °C and 37°C, respectively. **(A)** Impact on native saturated LysylPG species. **(B)** Impact on unsaturated LysylPG species, (n=5, ±S.D.).

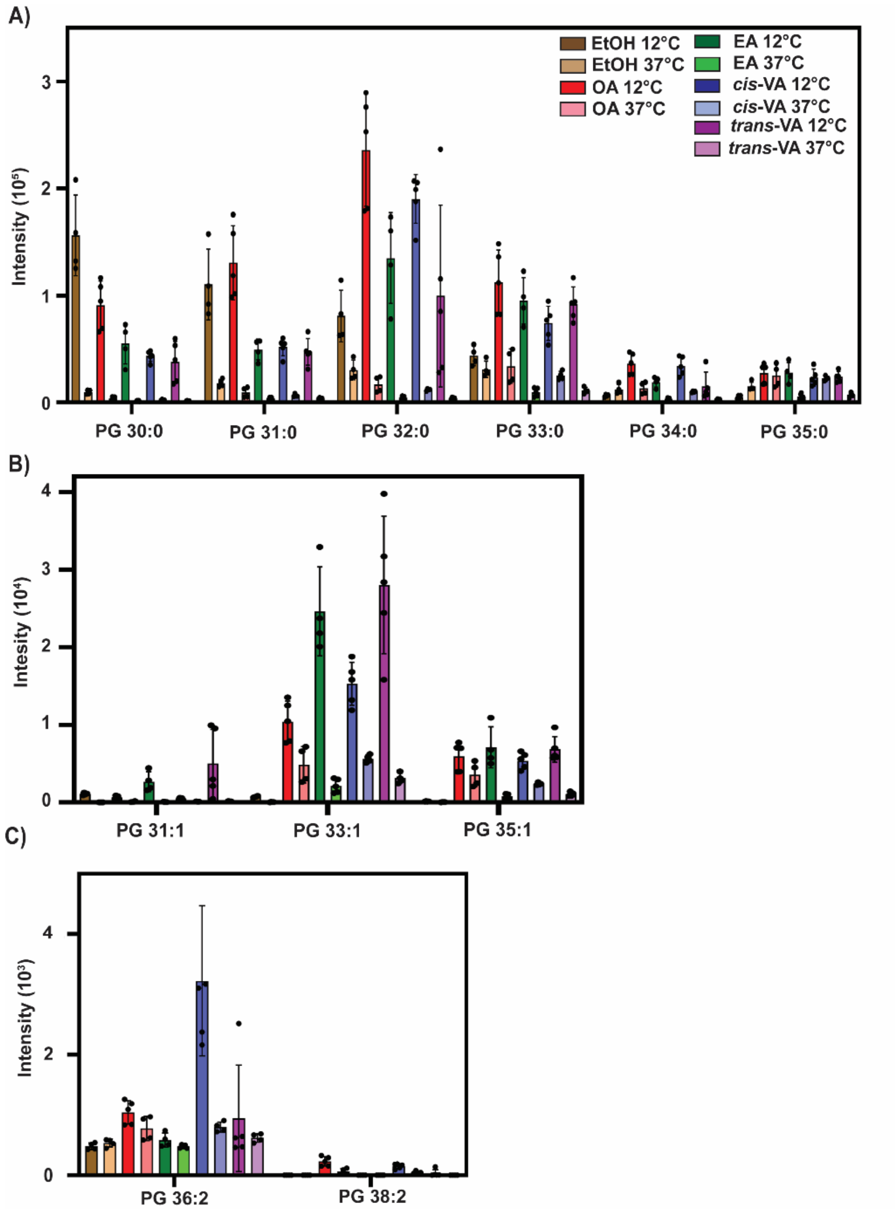


**SUPPLEMENTARY FIG S7:** Lipidomics analysis of PGs when supplemented with 100 μM of Oleic acid (9Z), Elaidic acid (9E), cis-vaccenic acid (11Z), or trans-vaccenic acid (11E) at 12°C and 37°C. (A) PGs containing only saturated fatty acids. (B) High-abundance PGs containing one unsaturated fatty acid. (C) Low-abundance PGs containing two unsaturated fatty acids, (n=5, ±S.D.).

**SUPPLEMENTARY TABLE S1:** Overview of growth stimulation of JE2 at 12 ˚C when supplemented with 10 μM and 100 μM of the indicated fatty acids and minimal inhibitory concentrations (MIC) reported in the literature (20) for growth at 37 ˚C.

| MONO-UNSATURATED FATTY ACIDS | | | |
|---|---|---|---|
| | 10 μM | 100 μM | MIC (μM) |
| Myristoleic Acid (C14:1Δ9) | + | + | 250 |
| Sapienic Acid (C16:1Δ6) | + | - | 30 |
| Palmitoleic Acid (C16:1Δ9) | + | - | 30 |
| Oleic Acid (C18:1Δ9) | + | +++ | >500 |
| Elaidic Acid (C18:1Δ9 *trans*) | + | ++ | - |
| *cis*-Vaccenic Acid (C18:1Δ11) | + | ++ | >500 |
| *trans*-vaccenic Acid (C18:1Δ11) | - | +++ | - |
| Erucic Acid (C22:1Δ13) | ++ | + | - |
| Nervonic Acid (C24:1Δ9) | ++ | + | - |

| Di-UNSATURATED FATTY ACIDS | | | |
|---|---|---|---|
| | 10 μM | 100 μM | MIC (μM) |
| Linoleic Acid (C18:2Δ9,12) | ++ | - | 250 |
| **Tri-UNSATURATED FATTY ACIDS** | | | |
| α-Linolenic Acid (C18:3Δ9,12,15) | + | - | 30 |
| γ-Linolenic Acid (C18:3Δ6,9,12) | ++ | - | 30 |
| **Tetra-UNSATURATED FATTY ACIDS** | | | |
| Stearidonic Acid (C18:4Δ6,9,12,15) | ++ | + | - |
| Arachidonic Acid (C20:4Δ5,8,11,14) | ++ | - | 30 |
| **Penta-UNSATURATED FATTY ACIDS** | | | |
| Eicosapentenoic Acid (C20:5Δ5,8,11,14,17) | ++ | - | - |
| **Hexa-UNSATURATED FATTY ACIDS** | | | |
| Docosahexenoic Acid (C22:6Δ 4,7,10,13,16,19) | ++ | - | - |

**SUPPLEMENTARY TABLE S2:** List of *S. aureus* strains used in this study

| Strain | Characteristics | Reference |
|---|---|---|
| JE2 | Derived from community-acquired MRSA strain USA300 | (Fey et al. 2013) |
| JE2 *fakA::Tn* | NR-46772 | (Fey et al. 2013) |
| JE2 *crtM::Tn* | Carotenoid-deficient mutant | (Braungardt and Singh 2019) |
| JE2 *crtM::Tn* +pCU*crtOPQMN* | *crtM* mutant of JE2 complemented with *crt* gene cluster in *trans* | (Braungardt and Singh 2019) |
| AH_LAC | Parent strain for the *geh* mutant | (Boles et al. 2010) |
| AH_LAC *Δgeh* | Lipase-deficient mutant | (Chen and Alonzo 2019) |
| JE2 *gtaB::Tn* | UTP-glucose-1-phosphate uridylyltranferase mutant | (Fey et al. 2013) |
| JE2*ΔugtP* | Diacylglycerol glucosyl transferase mutant | (Fey et al. 2013) |
